# Beyond years of schooling: Shifting genetic influences across educational milestones in two Norwegian cohorts

**DOI:** 10.1101/2025.10.08.680992

**Authors:** Eirik H. Kvalvik, Yunpeng Wang, Kristine B. Walhovd, Torkild H. Lyngstad, Ole Røgeberg

## Abstract

Although educational attainment is heritable, its conventional measurement in genetic research as years of education (EduYears) is not designed to reveal potential stage-specific genetic influences across discrete milestones. In two Norwegian cohorts (Norwegian Mother, Father and Child Cohort Study, N = 120,527; Norwegian Twin Registry, N = 8,910), we quantified the genetic contributions to completing high school, bachelor’s, master’s and PhD using genome-wide association studies (GWAS), polygenic indices (PGIs) and twin models. Transition-specific analyses, conditioning on prior success, revealed that observed-scale common-variant heritability (h^2^_SNP_) and PGI predictability followed an inverse-U pattern, peaking at the transition into higher education (h^2^_SNP_ ≈ 0.14; R^2^_Tjur_ ≈ 0.05) before declining for postgraduate degrees. Genetic correlations (r_g_) with large-scale GWAS of EduYears (EA4) and intelligence (IQ3) were high for early transitions but declined markedly for later ones (e.g., r_g_ with EA4 from ≈ 0.92 to ≈ 0.38). In cumulative analyses, aggregating liability across prior milestones, the gap between twin- and SNP-based heritability narrowed at higher levels of attainment (h^2^_twin_ ≈ 0.6→0.3; h^2^_SNP_ ≈ 0.22→0.19), while the genetic overlap between distant milestones diminished (r_g_ ≈ 0.92→0.71). These patterns, obscured by EduYears metrics, highlight a dynamic genetic architecture across educational milestones, refining polygenic prediction and addressing misconceptions about uniform genetic influences on educational progression.

## Introduction

Educational attainment (EA) is a well-established predictor of adult health^1,2^, economic productivity^3^ and overall well-being^4^. It therefore features prominently in social science research, both as an outcome, a control variable, and an explanatory factor. In recent years it has also become one of the most widely studied social phenotypes in behavioural genetics. Twin and molecular studies show that the customary measure, years of education (often termed “EduYears” in the literature), is substantially heritable^5–7^. Yet a simple year-count overlooks a basic structural fact: progress depends on clearing discrete milestones, such as finishing high school, earning a bachelor’s degree, completing a master’s, or obtaining a doctorate, each with its own demands. Major demographic surveys reflect this categorical reality by asking, “What is the highest degree or level of school you have completed?” rather than “How many years did you attend school?”* Even researchers using EduYears implicitly acknowledge this milestone-based structure, as they typically assign “normed years” based on credentials earned rather than actual time spent in the educational system. This structure has long been the explicit focus of sociological models of stratification, which treat schooling as a sequence of decision points^10–12^.

While these models have mapped social and institutional influences on progression, they have paid little attention to whether, and to what extent, genetic influences differ from one milestone to the next. Such differences are plausible, as the educational journey likely demands a shifting portfolio of cognitive and non-cognitive skills^13–15^, is shaped by complex gene-environment interactions^16^, and involves selection processes that progressively narrow observed variance among those who advance, which can attenuate associations within selected groups^17^. This issue has gained urgency as access to higher education expands globally and as technology increases demand for high-skill work^18,19^. The tools used in genetic research, however, have not fully kept pace. Molecular efforts have culminated in progressively larger genome-wide association studies (GWAS) of EduYears, collectively termed EA1 through EA4^20–23^. The polygenic indices (PGIs) derived from these studies have relied on this continuous metric, without accounting for the discrete, hierarchical nature of educational progression. This may lead to misconceptions about the uniformity of genetic influences across the educational ladder and leaves the utility of these PGIs for discriminating success in higher education uncertain^24^.

We address this gap by leveraging the Norwegian Mother, Father and Child Cohort Study (MoBa; 120,527 genotyped adults) and the Norwegian Twin Registry (NTR; 8,910 twins).

Across four educational milestones, we characterize genetic patterns using two complementary frameworks.

First, using a transition-specific framework that conditions on prior success, we isolate the genetic factors relevant to each sequential step. After conducting a GWAS for each milestone transition, we examine external genetic correlation (r_g_) with large-scale meta-analyses of EduYears (EA4^23^) and intelligence (IQ3^25^); estimate their single nucleotide polymorphism (SNP) heritability (h^2^_SNP_); and test the predictive power of an EduYears-based PGI (EA4-derived).

Second, using a cumulative framework, we model the total genetic liability required to achieve each milestone status. This approach enables us to assess the genetic overlap between the milestones themselves and to contrast common-variant heritability (h^2^_SNP_) with the full additive genetic contribution from biometric twin models (h^2^_twin_), which were only feasible under this framework.

Our findings show that analysing credentials instead of years uncovers patterns invisible to a purely time-based approach and refines the interpretation of PGIs now widely used in sociogenomic research. Rather than replacing years-based approaches, we demonstrate the substantial information gained by recognizing education’s milestone-based structure, where success is measured by credentials earned rather than time invested.

## Results

To address the limitations of treating EA as a continuous measure, we reconceptualized it as a sequence of discrete milestones. Figure 1 presents the conceptual frameworks guiding our study.

**Figure 1.**
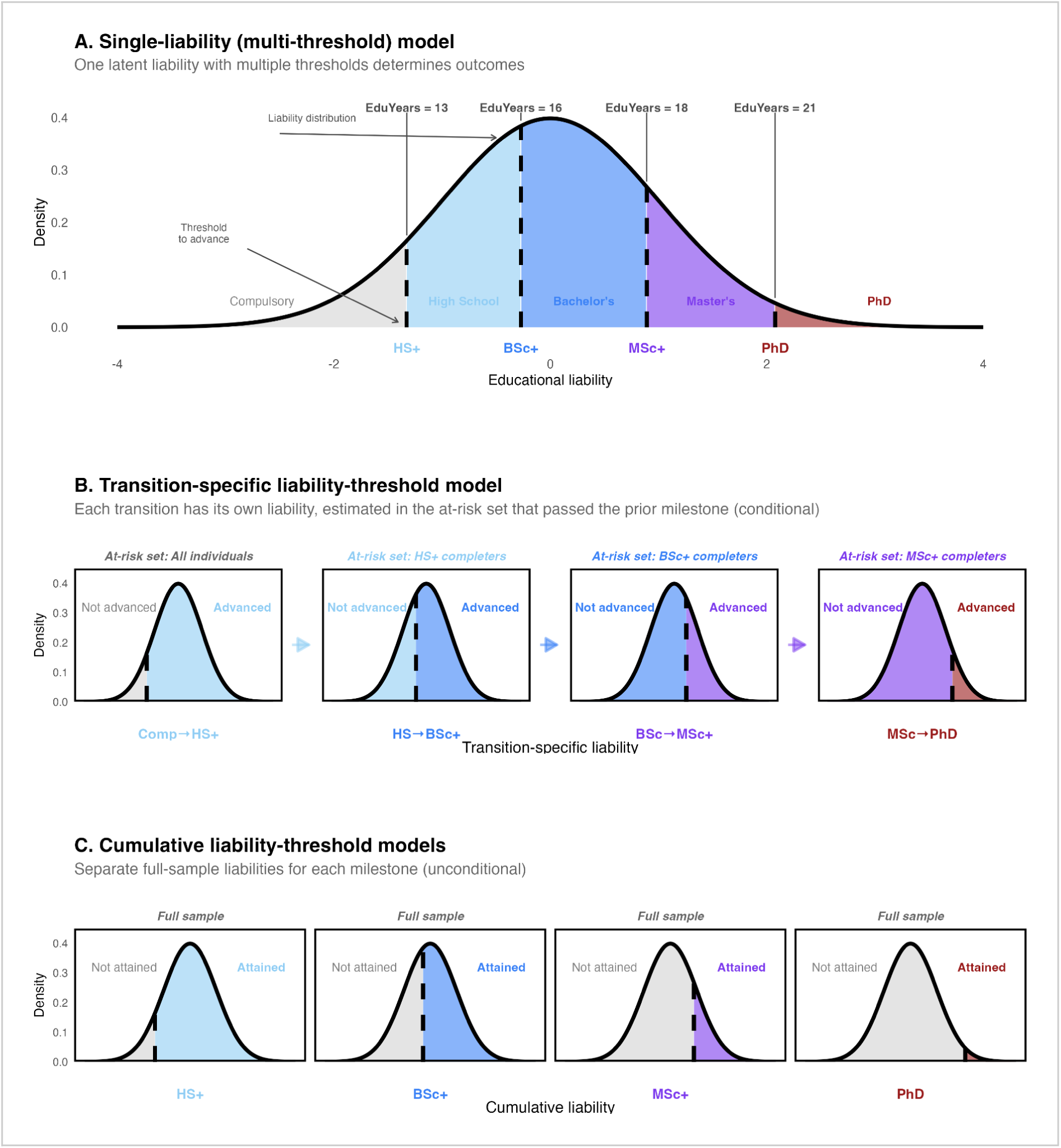
Conceptual models of genetic influence on educational attainment. **A, Single liability threshold model.** This assumes qualitative uniformity, where the same genetic factors influence the entire educational journey. However, it allows for quantitative non-linearity via uneven thresholds. This depiction is more nuanced than conventional EduYears GWAS, which typically impose both qualitative and quantitative (linear) uniformity. **B, Transition-specific liability threshold model.** In contrast, this framework models each milestone transition as governed by its own liability distribution, estimated conditionally in the at-risk population that cleared the prior milestone (color-coded ‘at-risk’ sets). This allows for both qualitative differences (different genes being important) and quantitative non-linearity (varying genetic effects) across the educational journey. **C, Cumulative liability threshold models.** This approach models the aggregated genetic liability for attaining each milestone (at least that level vs. below) by estimating separate liability distributions unconditionally in the full population.

Panel A illustrates the traditional single-liability model, which assumes qualitative uniformity, where the same genetic factors influence the entire educational journey. However, it allows for quantitative non-linearity via uneven thresholds. This depiction is more structured than conventional EduYears GWASs, which typically impose both qualitative and quantitative (linear) uniformity by assuming equal, additive effects per year of schooling on the observed scale. In contrast, our work employs two complementary frameworks.

Panel B depicts our transition-specific liability-threshold model, where each milestone transition is governed by a distinct liability distribution (allowing for both qualitative heterogeneity and quantitative variation) conditional on prior success. We applied this framework to investigate the genetic basis of four key educational milestone transitions.

Panel C shows the cumulative liability models. In contrast to the conditional approach in Panel B, these models are estimated unconditionally in the full sample. By estimating a distinct liability distribution for each milestone, they capture the aggregated genetic liability for achieving each milestone status by comparing everyone who attained a specific level or higher against everyone who did not. This framework is essential for assessing the total genetic overlap between distant milestones and for enabling analyses like twin modeling, which benefit from using the entire population.

This dual approach allows us to disentangle the genetics of the process of educational progression from the genetics of the final outcome achieved. We denote “at least high school” as HS+, “at least bachelor’s” as BSc+, “at least master’s” as MSc+, and PhD as doctoral completion; Comp indicates compulsory education or below. A milestone transition is written X→Y, with cases at level Y and controls at X only (e.g., HS→BSc+ compares BSc+ to HS-only). We first present the findings from the transition-specific analyses.

### Transition-specific genome-wide association analyses of educational milestones

Following our transition-specific framework (Figure 1B), we conducted genome-wide association analyses in the MoBa genotyped adults (N = 120,527). Specifically, we modeled four sequential transitions: Comp→HS+, HS→BSc+, BSc→MSc+, and MSc→PhD. For each analysis, cases were individuals who completed the later milestone, and controls were those who achieved only the immediately preceding level. This approach isolates the genetic factors specific to each transition, conditional on prior success, rather than those accumulated across all educational milestones. After standard quality-control (QC) filtering (e.g., imputation quality, minor allele frequency, Hardy–Weinberg equilibrium), we fit a linear mixed model (fastGWA) with covariates (sex, birth-cohort bins, genotyping batch, PC1–PC10). Genome-wide significance was set at two-sided *p* < 5 × 10^−8^. Calibration from LD score regression (LDSC) was acceptable across milestone transitions (intercepts ≈ 1.00–1.06; attenuation ratios ≈ 0.12–0.17; Supplementary Table S1), indicating that most inflation reflects polygenicity rather than residual stratification^26^. These transition-specific GWASs provided the inputs for the subsequent crosstrait LDSC r_g_ and h²_SNP_ analyses. We report observed-scale h²_SNP_ estimates for transition phenotypes to avoid assumptions about conditional population prevalences; r_g_ estimates are scale-free and unaffected by this choice.

### Genetic correlations: early transitions align with educational attainment and intelligence, but overlap attenuates at the postgraduate stage

Building on these GWASs, cross-trait LDSC r_g_ estimates showed that the genetic factors captured by our transition-specific analyses aligned closely with those underlying broad measures of schooling (EA4) and general cognitive ability (GCA; IQ3) up to the entry into higher education, but diverged at the postgraduate stage (Fig. 2).

**Figure 2.**
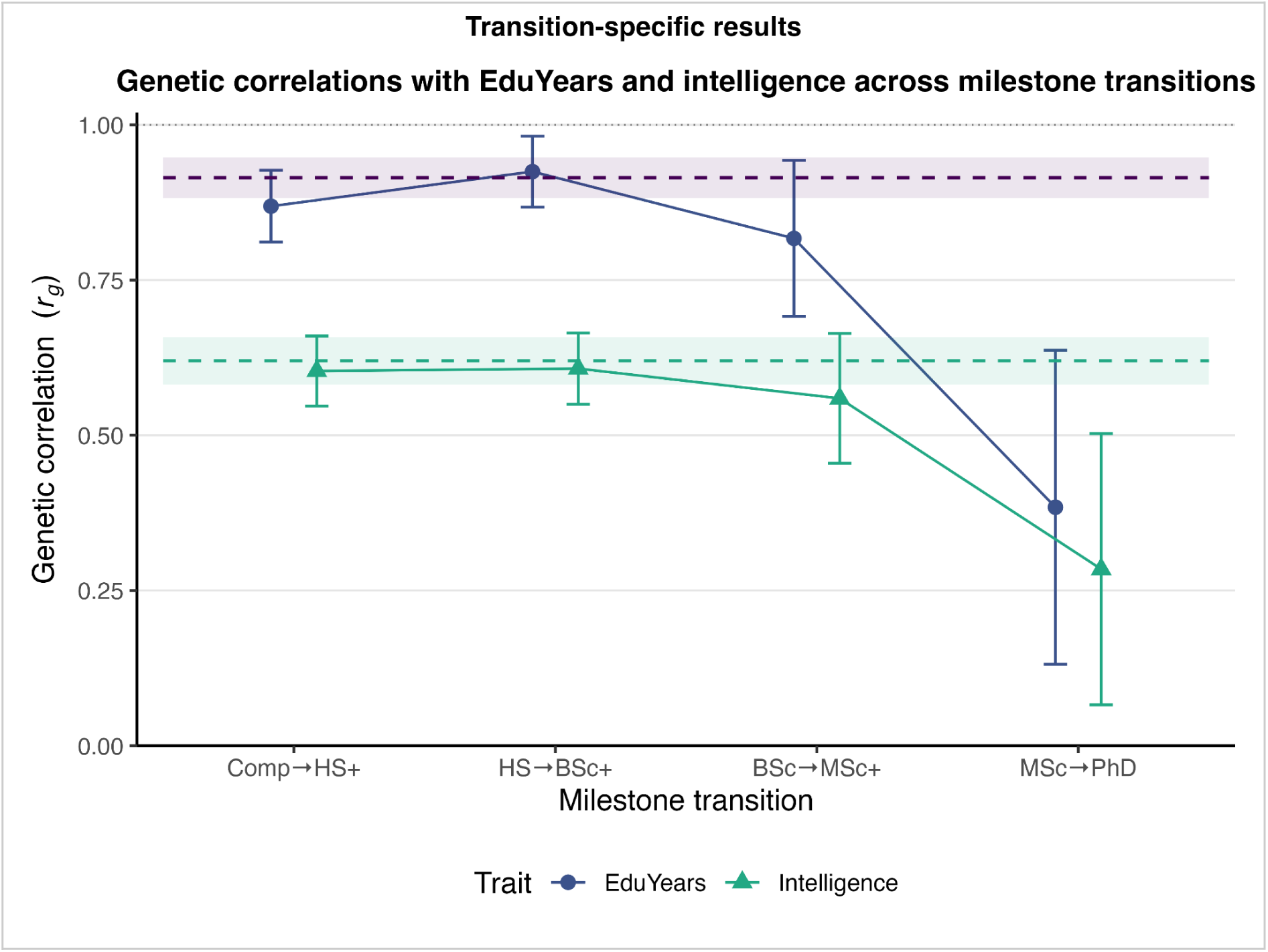
Genetic correlations between transition-specific GWASs in MoBa and external GWAS meta-analyses of EduYears and intelligence. LDSC genetic correlations (r_g_) between each transition-specific GWAS (Comp→HS+, HS→BSc+, BSc→MSc+, MSc→PhD) and external GWAS meta-analyses of EduYears (EA4^23^; navy circles) and intelligence (IQ3^25^; teal triangles). Error bars indicate 95% confidence intervals (CIs). Horizontal dashed lines with shaded 95% CI ribbons represent benchmark correlations from a GWAS of EduYears within the MoBa cohort, showing its correlation with EA4 (purple) and IQ3 (teal). The grey dashed line indicates r_g_ = 1.0 for reference. The plot shows that the genetic overlap with both EduYears and intelligence is strong for early milestone transitions but declines markedly at the postgraduate stage, particularly for MSc→PhD.

To contextualize our findings, we first established reference benchmarks using our MoBa cohort, serving as empirical ceilings for genetic overlap with EA4 and IQ3. The r_g_ between MoBa EduYears and EA4 indicated near-complete overlap (≈ 0.93, 95% CI [0.90–0.96]). Similarly, the r_g_ between MoBa EduYears and IQ3 established the expected relationship between EA and GCA (≈ 0.63, [0.58–0.67]).

Against these benchmarks, early milestone transitions showed high genetic overlap with EduYears (Comp→HS+: r_g_ ≈ 0.87 [0.81–0.93]; HS→BSc+: r_g_ ≈ 0.92 [0.87–0.98]) and with GCA (Comp→HS+: r_g_ ≈ 0.60 [0.55–0.66]; HS→BSc+: r_g_ ≈ 0.61 [0.55–0.66]), indicating that they essentially captured the same common-variant architecture found in EA4 and IQ3. The pattern shifted at postgraduate levels. BSc→MSc+ showed diminished correlations with EduYears (r_g_ ≈ 0.82 [0.69–0.94]) and GCA (r_g_ ≈ 0.56 [0.45–0.66]), though these remained substantial. MSc→PhD marked the sharpest decline, with values far below our benchmarks for EduYears (r_g_ ≈ 0.38 [0.13–0.64]) and GCA (r_g_ ≈ 0.28 [0.07–0.50]), indicating reduced alignment with the genetic influences captured by EA4 and IQ3, though with wider confidence intervals at this transition.

### Observed-scale SNP heritability follows an inverse-U across milestone transitions

Following the patterns of genetic overlap, the magnitude of the common-variant signal also varied across the milestone transitions, with observed-scale (see Methods) LDSC estimates of SNP-based heritability (h²_SNP_) following an inverse-U pattern (Figure 3a): it was modest for the move from compulsory schooling to high school (Comp→HS+: h²_SNP_ ≈ 0.07 [0.06–0.09]), peaked for high-school completers progressing to a bachelor’s (HS→BSc+: h²_SNP_ ≈ 0.14 [0.12–0.16]), and declined at postgraduate stages (BSc→MSc+: h²_SNP_ ≈ 0.1 [0.07–0.12]; MSc→PhD: h²_SNP_ ≈ 0.07 [0.01–0.13]), though with a much wider CI for PhD reflecting its smaller effective sample size (*N*_eff_ = 8,185; see Table 1).

**Figure 3.**
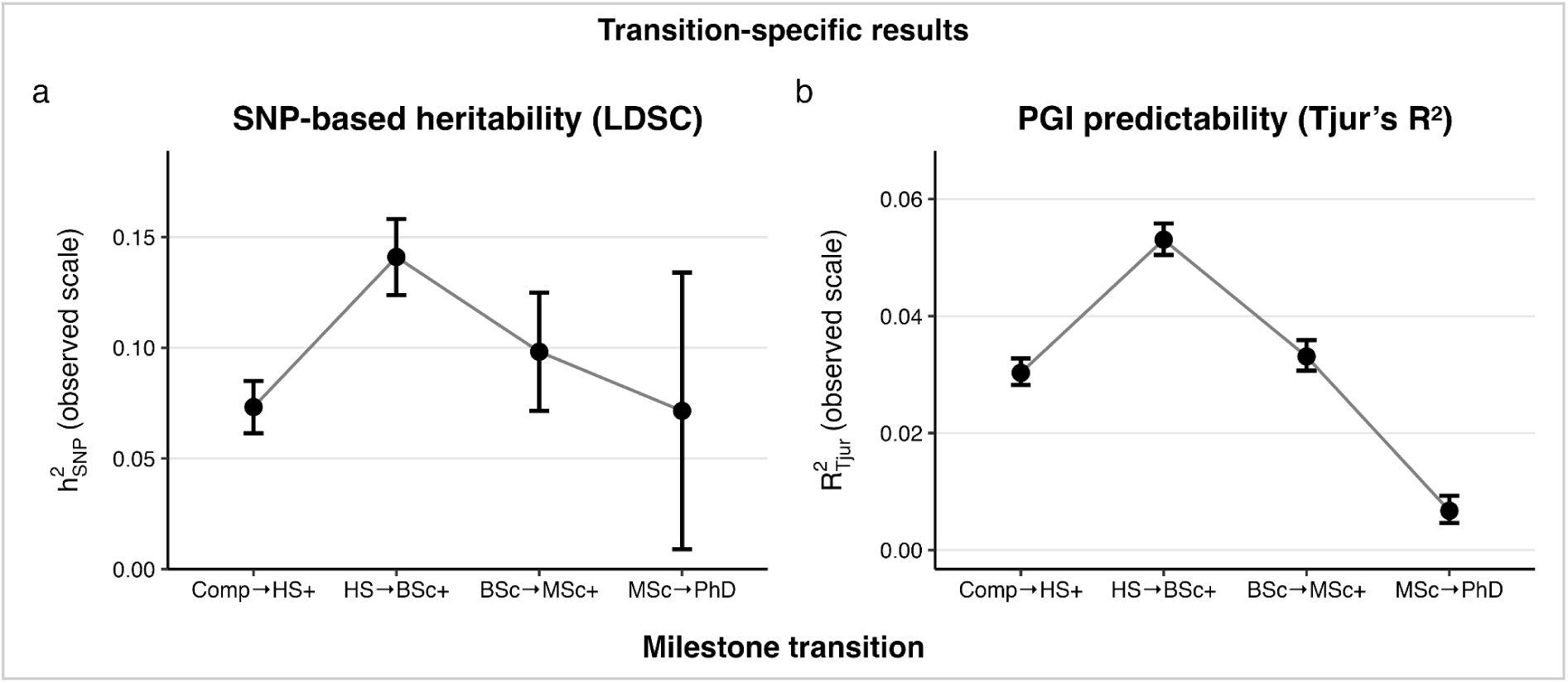
Common-variant signal and PGI discrimination at successive educational transitions. **a,** Observed-scale LDSC estimates of SNP-based heritability (h²_SNP_) for each transition: Comp→HS+, HS→BSc+, BSc→MSc+, MSc→PhD. Points show estimates; vertical bars show 95% CIs. Heritability peaks at HS→BSc+ and declines at postgraduate stages. **b,** Predictive performance of an EA4-derived polygenic index (PGI) for each transition, summarized by Tjur’s R^2^ from adjusted probit models; 95% bootstrap (1,000 resamples) CIs shown. Discrimination follows an inverse-U, with a marked drop at MSc→PhD, consistent with increasing selection/range restriction in the at-risk set. Both panels are on the observed scale; no liability transformation is applied to transition outcomes.

**Table 1.** Sample sizes and effective sample sizes (*N*_eff_) for educational milestone analyses. The “*N* (Total)” column lists the full cohort size for cumulative outcomes and the at-risk set (those who cleared the prior milestone) for transition-specific outcomes. Effective sample size (*N*_eff_), a key indicator of statistical power for binary traits, is calculated as *N*_eff_ = 4/(1/*N*_cases_+1/*N*_controls_) = 4*NP*(1−*P*) ^39^, where *P* is the case proportion. Because power scales with *N*_eff_, information is maximized near 50:50 case–control balance and declines as prevalence departs from 0.5. For the first milestone, cumulative (HS+) and transition-specific (Comp→HS+) definitions coincide. In MoBa, cumulative definitions yield larger *N*_eff_ than transition–specific ones at BSc+, MSc+, and PhD (≈ +18.7%, +17.4%, +9.2%), while NTR PhD has a very small *N*_eff_ (313), implying wide confidence intervals.

| Cohort | Approach | Milestone | *N*<br>(Cases) | *N*<br>(Controls) | *N*<br>(Total) | *P*<br>(Case<br>proportion) | *N*<sub>eff</sub><br>(Effective<br>Sample Size) |
| --- | --- | --- | --- | --- | --- | --- | --- |
| NTR | Cumulative | HS+ | 7,782 | 1,128 | 8,910 | 87.3% | 3,941 |
|  |  | BSc+ | 4,770 | 4,140 | 8,910 | 53.5% | 8,865 |
|  |  | MSc+ | 1,470 | 7,440 | 8,910 | 16.5% | 4,910 |
|  |  | PhD | 79 | 8,831 | 8,910 | 0.9% | 313 |
| MoBa | Cumulative | HS+ | 109,398 | 11,129 | 120,527 | 90.8% | 40,406 |
|  |  | BSc+ | 73,188 | 47,339 | 120,527 | 60.7% | 114,983 |
|  |  | MSc+ | 22,464 | 98,063 | 120,527 | 18.6% | 73,109 |
|  |  | PhD | 2,277 | 118,250 | 120,527 | 1.9% | 8,936 |
| MoBa | Transition<br>-specific | Comp→HS+ | 109,398 | 11,129 | 120,527 | 90.8% | 40,406 |
|  |  | HS→BSc+ | 73,188 | 36,210 | 109,398 | 66.9% | 96,899 |
|  |  | BSc→MSc+ | 22,464 | 50,724 | 73,188 | 30.7% | 62,276 |
|  |  | MSc→PhD | 2,277 | 20,187 | 22,464 | 10.1% | 8,185 |

### Polygenic indices stratify progression up the educational ladder, while its predictive power peaks at the transition into higher education

Extending these heritability insights to prediction, we evaluated how the EA4 PGI forecasts each milestone transition in MoBa (Figure 3b). Using probit regression adjusted for sex, birth cohort, and ten ancestry principal components, we quantified predictive power using Tjur’s coefficient of discrimination (R^2^_Tjur_) on the observed scale. The results followed an inverse-U pattern: predictability was moderate for Comp→HS+ (R^2^_Tjur_ ≈ 0.03), peaked for HS→BSc+ (R^2^_Tjur_ ≈ 0.05), and then waned for the postgraduate transitions (BSc→MSc+ ≈ 0.03; MSc→PhD < 0.01).

Figure 4 provides further insight by contrasting the mean PGI levels between advancing and non-advancing groups at each milestone. This reveals a decoupling between mean-level separation and the PGI’s predictive power shown in Figure 3b. While the mean PGI was consistently higher for the group that progressed at every milestone (Figure 4a), the distributions of PGI scores for advancing and non-advancing groups showed extensive overlap, which was particularly evident for the PhD transition (Figure 4b). This highlights that although the PGI effectively stratifies the educational hierarchy on average, its ability to discriminate between individuals who will or will not advance to the next stage is limited, particularly at boundary milestones where base rates are very high (Comp→HS+) or very low (MSc→PhD).

**Figure 4.**
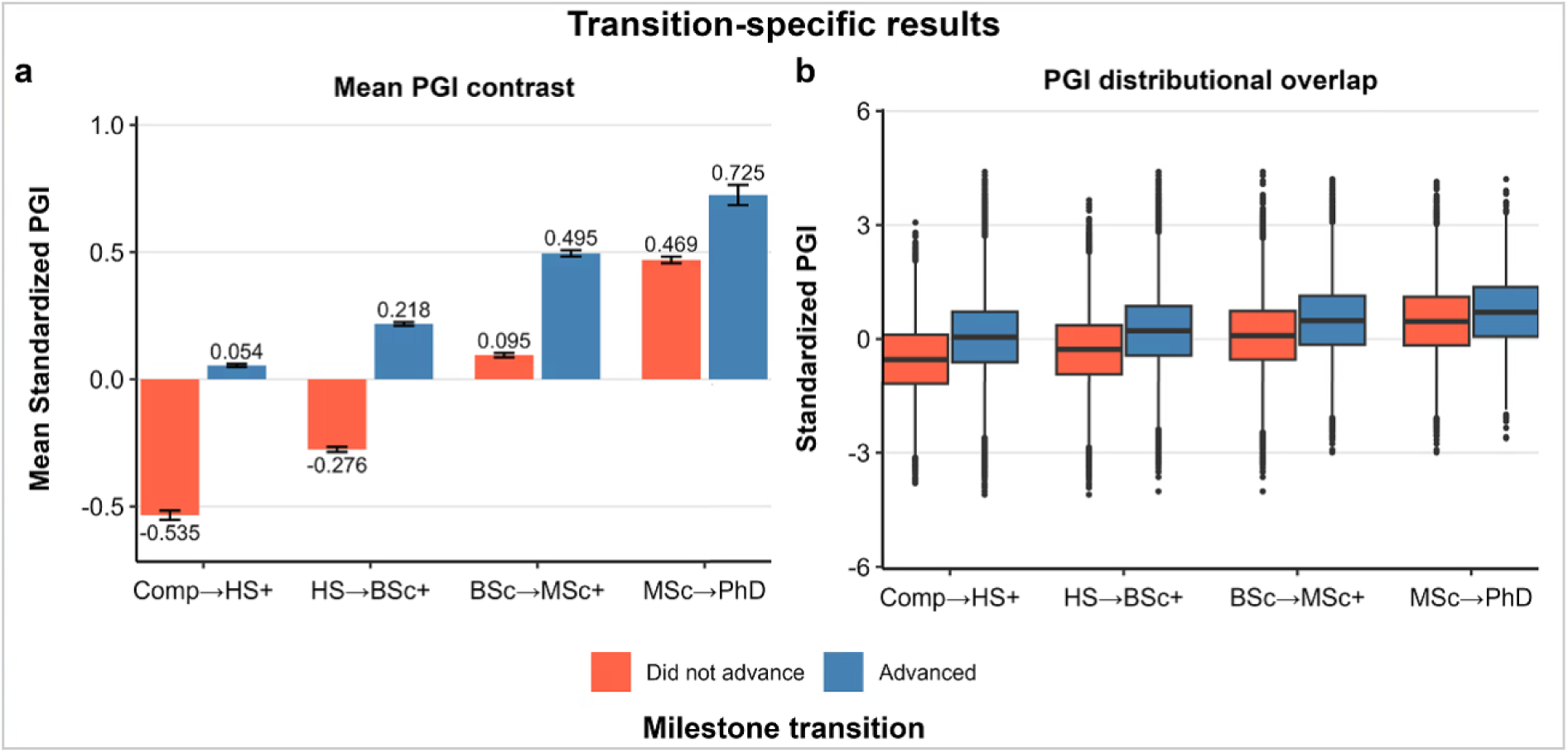
PGI differences at each educational milestone transition. **a**, **Mean PGI contrast.** Bars show the mean standardized PGI (z-score; overall MoBa mean = 0, s.d. = 1) for individuals who did not advance (orange) and those who advanced (blue) to each milestone. Error bars represent ± 1.96 s.e.; numeric labels indicate the group means. These data illustrate two main patterns: advancing groups consistently carry higher mean PGIs than non-advancing groups at each milestone, and this mean difference (gap) narrows progressively across subsequent transitions (from ≈ 0.58 s.d. at HS+ to ≈ 0.25 s.d. at PhD) **b**, **PGI distributional overlap**. Box plots show the full PGI distribution for the same groups, illustrating the considerable overlap in scores between advancing and non-advancing groups for each milestone transition.

### Cumulative genetic influences: Inter-milestone overlap and heritability comparisons

To complement the transition analyses, we re-estimated each milestone under a cumulative definition (Fig. 1C), quantifying the total genetic liability required to reach at least each level (HS+, BSc+, MSc+, PhD) by contrasting it with all lower levels in the full sample. This design was essential for three reasons. First, it enables within-cohort bivariate genetic correlations among milestones via bivariate GREML (identifiable on the full sample). Second, it allows a twin–molecular comparison by reporting common-variant and twin-based heritabilities on the same (liability) scale. Third, it provides two complementary estimators of SNP heritability for robustness: LDSC from summary data and BOLT-REML from individual-level data. The GWAS pipeline matched the transition analyses (QC, fastGWA LMM, covariates, threshold). LDSC calibration was again acceptable (intercepts ≈ 1.02–1.08; ratios ≈ 0.09–0.16; Supplementary Table S1).

Using bivariate BOLT-REML, genetic correlations (Fig. 5) were highest for adjacent steps (e.g., HS+ vs BSc+, r_g_ ≈ 0.92) and declined with educational distance (e.g., HS+ vs PhD, r_g_ ≈ 0.71). This gradient indicates substantial shared liability across neighboring milestones together with increasing stage-specific influences as milestones become more distant.

**Figure 5.**
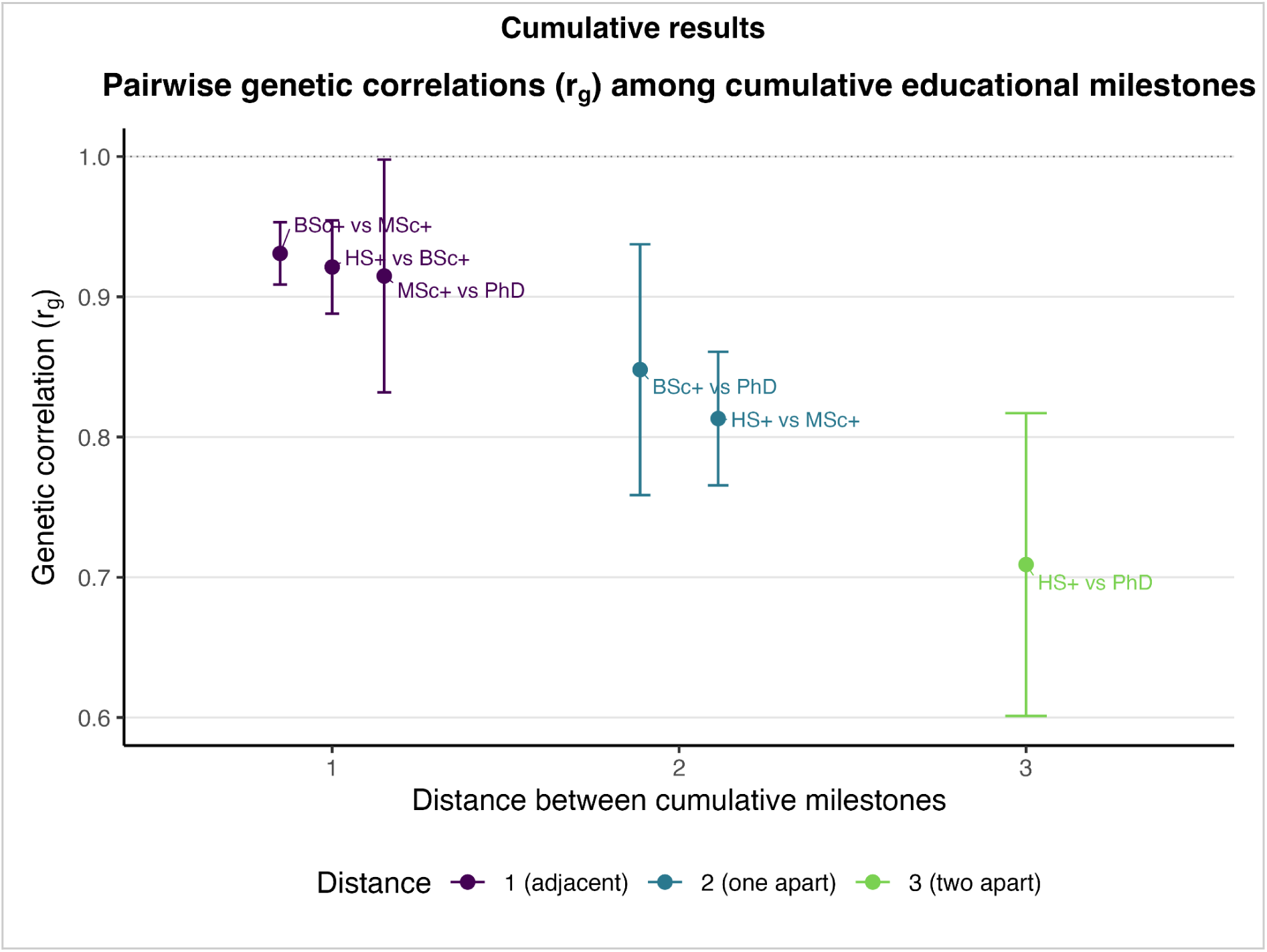
Pairwise genetic correlations (r_g_) among cumulative educational milestones in MoBa. Genetic correlations (r_g_) were estimated with bivariate BOLT-REML for all pairs of cumulative milestones (HS+, BSc+, MSc+, and PhD). Points show r_g_ estimates with 95% confidence intervals. Color encodes the number of steps separating each pair and matches the x-axis “distance”: 1 = adjacent (HS+ vs BSc+, BSc+ vs MSc+, MSc+ vs PhD), 2 = one step apart (HS+ vs MSc+, BSc+ vs PhD), and 3 = two steps apart (HS+ vs PhD). The dashed horizontal line marks r_g_ = 1. Genetic overlap is highest for adjacent milestones and declines with increasing distance, with the lowest r_g_ for HS+ vs PhD and the widest uncertainty for comparisons that include PhD.

### Twin–SNP heritability gap peaks at high-school completion and narrows at higher degrees

To quantify the full additive genetic contribution to each cumulative milestone, we analyzed data from 8,910 twins in the Norwegian Twin Registry. Additive twin heritability (h^2^_twin_) from Bayesian liability-threshold ACE models (Figure 6a) was highest for HS+ (≈ 0.60, 95% credible interval [0.39, 0.76]) and declined progressively for BSc+ (≈ 0.49, [0.36, 0.64]), MSc+ (≈ 0.42, [0.23, 0.61]), and PhD (≈ 0.30, [0.02, 0.62]). Although intervals widen at rarer milestones, the downward trend in point estimates is clear. Environmental components show contrasting patterns: the shared environment (C) follows an inverse-U shape, peaking around BSc+ and MSc+ (both ≈ 0.31), whereas the unique environment (E), which also includes measurement error, shows a U-shape, lowest at BSc+ (≈ 0.20) and highest at PhD (≈ 0.47). Full ACE results are provided in Supplementary Table S3 and Figure S26.

**Figure 6.**
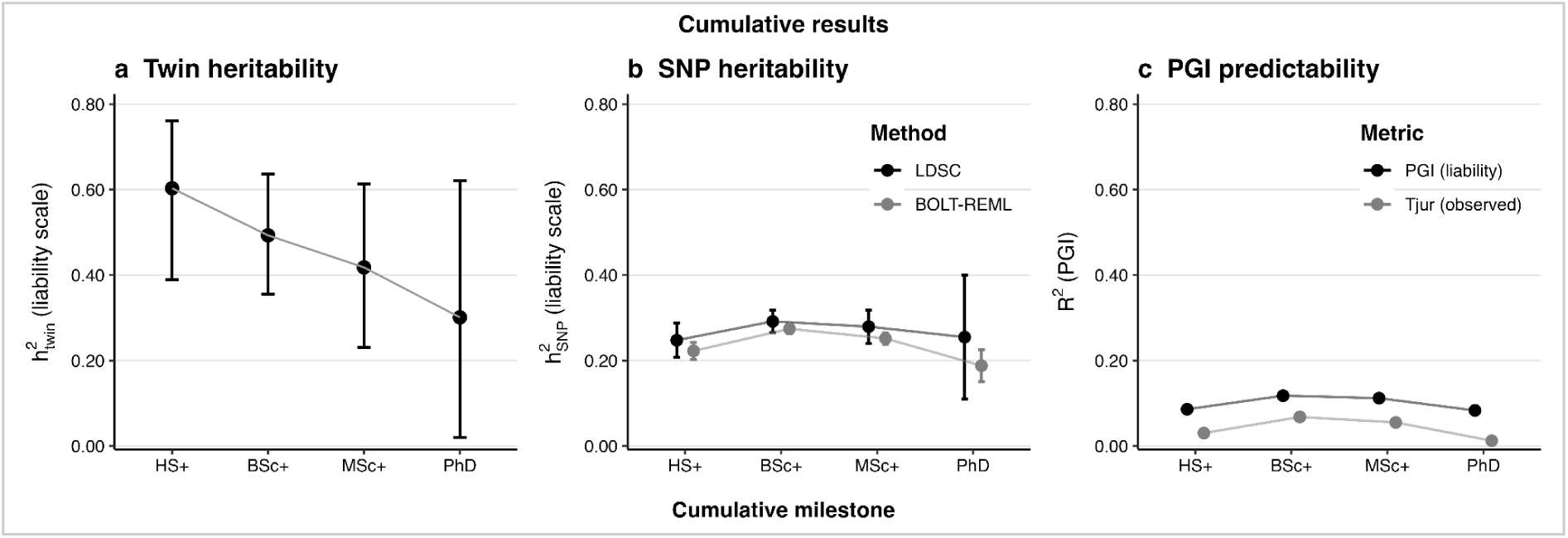
Cumulative genetic influences across educational milestones. **a, Twin heritability.** Additive twin heritability (h^2^_twin_) estimated with Bayesian liability-threshold ACE models from the Norwegian Twin Registry; points are posterior means and whiskers are 95% credible intervals (liability scale). **b, SNP heritability.** Common-variant heritability (h^2^_SNP_) estimated by LDSC (black) and BOLT-REML (grey); whiskers are 95% confidence intervals (liability scale). **c, PGI predictability.** Two complementary metrics for the EA4-based polygenic index. PGI (liability) is the variance on the liability scale explained by the PGI derived from the probit slope with a standardized residualized PGI following Lee et al.^27^. Tjur (observed) is Tjur’s coefficient of discrimination (R^2^_Tjur_) from probit models on the observed scale. Whiskers are 95% bootstrap CIs (1,000 resamples) for both metrics. Across panels, h^2^_twin_ declines from HS+ to PhD, h^2^_SNP_ and PGI performance rises from HS+ to BSc+ and attenuate at advanced degrees.

Using identical cumulative contrasts in MoBa, we then estimated common-variant SNP heritability (h^2^_SNP_) with two orthogonal methods, summary-level LDSC and individual-level BOLT-REML (Figure 6b). LDSC is robust to many forms of stratification via its intercept but is typically less statistically efficient (wider standard errors) than individual-level REML; agreement between the two increases confidence in the estimates. To permit direct comparison with twin-based heritability (h^2^_twin_), observed-scale estimates were converted to the liability scale (see Methods). Both methods show a similar inverse-U pattern: values increase from HS+ (≈ 0.25 LDSC; ≈ 0.22 BOLT-REML) to a peak at BSc+ (≈ 0.29; ≈ 0.27), then decline for MSc+ (≈ 0.28; ≈ 0.25) and PhD (≈ 0.25; ≈ 0.19). To enable a direct comparison with the twin-based results, these SNP-based heritabilities are presented on the liability scale.

Finally, we assessed the predictability of an EA4-based PGI for the cumulative milestones using two complementary metrics (Figure 6c). First, we report the variance explained on the liability scale, which enables a direct comparison with our heritability estimates, derived from the probit slope with a standardized residualized PGI following Lee et al.^27^: HS+ ≈ 0.09, BSc+ ≈ 0.12, MSc+ ≈ 0.11, and PhD ≈ 0.08. Second, we report Tjur’s R², which captures discrimination on the observed scale: HS+ ≈ 0.03, BSc+ ≈ 0.07, MSc+ ≈ 0.06, and PhD ≈ 0.01. Both metrics peak near the bachelor’s level and attenuate at MSc+ and PhD.

Comparing these estimates reveals distinct patterns across methods (Figure 6). Twin heritability decreases monotonically from HS+ to PhD, while both SNP heritability and PGI predictability follow an inverse-U pattern, peaking at BSc+. The gap between twin and SNP heritability estimates is largest at HS+ and narrows for subsequent milestones.

## Discussion

Educational attainment is not genetically uniform across the schooling sequence. Treating education as both a series of sequential transitions and a set of cumulative statuses reveals two decisive properties of its dynamic genetic architecture: qualitative (compositional) heterogeneity, meaning the mix of genetic influences is not the same at every stage, and quantitative (magnitude) heterogeneity, meaning the overall strength of genetic influence changes across stages.

Evidence for qualitative heterogeneity comes from three patterns. First, in the transition framework, genetic correlations with broad factors, EduYears (EA4) and general cognitive ability (IQ3), are high at early transitions but attenuate at BSc→MSc+ and MSc→PhD, indicating that postgraduate progression is less aligned with these general dimensions. Second, in the cumulative framework, internal pairwise genetic correlations decline with educational distance, whereas each cumulative milestone retains a high and stable correlation with EA4 and IQ3 through MSc+ (Supplementary Fig. S25). This contrast points to a two-component architecture: a shared general component that persists across most milestones and stage-specific components that increasingly differentiate higher levels. Third, the gap between twin- and SNP-based heritability is largest at HS+ and narrows thereafter, implying that the fraction of heritable variance captured by common SNPs, relative to total additive variance, is not constant across milestones. Together, these lines of evidence indicate qualitative heterogeneity rather than a single, stage-invariant architecture.

Evidence for quantitative heterogeneity is seen in the fact that the magnitude of genetic influence varies by milestone, with the transition framework showing a sharper inverse-U and the cumulative framework a flatter one. On the observed scale for transition outcomes, both SNP-based heritability and PGI discrimination (Tjur’s R^2^) are modest for the first step (Comp→HS+), peak at entry into higher education (HS→BSc+), and decline at postgraduate transitions. For cumulative outcomes on the liability scale, SNP-based heritability and the PGI (liability) are highest around BSc+ and lower at HS+ and PhD, whereas twin-based liability heritability declines monotonically from HS+ to PhD. Because transition metrics are on the observed scale and cumulative metrics are on the liability scale, we compare shapes within each framework and avoid cross-scale comparisons of absolute magnitudes. Collectively, these results show that the amount of heritable variation expressed at each milestone is not static but changes across the educational sequence. We next consider mechanisms that could generate these patterns, contrasting measurement limits in external comparators with selection coupled with base-rate constraints in the at-risk sets (case fractions far from 0.5 that reduce observed-scale variance and predictive discrimination), and we argue that the within-MoBa shapes are most parsimoniously explained by the latter.

Measurement limits in our primary comparator, the EA4 meta-analysis of EduYears^23^, matter chiefly for the absolute level of our external cross-trait genetic correlations and PGI performance, rather than for the within-MoBa shapes. Our cross-trait analyses rely on an EA4 subset that excludes 23andMe, relies primarily on UK Biobank’s binary “any college or university” item, and codes credentials via ISCED-1997^28^, which collapses bachelor and master degrees (see Supplementary Note 1). These choices compress distinctions that our design preserves and can attenuate overlap precisely where those distinctions matter. At the MSc→PhD step, the external genetic correlation drops more for EA4 than for IQ3; this pattern is compatible with additional attenuation from credential collapsing and is also consistent with a shift in trait composition, for example a changing balance of cognitive and non-cognitive influences at the PhD stage. Accordingly, we view EA4’s measurement limits as important for interpreting magnitudes in external benchmarks, but as shown next, not the principal driver of the inverse-U patterns observed within MoBa.

Selection within the at-risk sets provides the primary account of the transition patterns. At the first step, high-school completion is near-universal, so non-completers form a small subgroup drawn from the lower tail of the EA-related polygenic distribution; at the last step, PhD completers form a small subgroup drawn from the upper tail. In both settings, selection with unbalanced case fractions (far from 0.5) compresses observed-scale signal in the at-risk sets, producing an inverse-U in transition-specific observed-scale h^2^_SNP_ and PGI discrimination (Tjur’s R^2^). The PGI is EA4-derived and thus inherits the credential collapsing of the EduYears coding; despite this, its transition pattern in MoBa closely mirrors that of milestone-GWAS-based h^2^_SNP_, which is not subject to those measurement choices. This convergence indicates that the within-MoBa inverse-U is unlikely to be driven primarily by EA4’s measurement limits and is instead best explained by selection with unbalanced base rates in the at-risk sets. Under the transition framework, repeated conditioning progressively narrows variance and accentuates this shape; under the cumulative framework, which is estimated in the full cohort, the effect is diluted, yielding flatter estimates across milestones.

For the cumulative milestones, the gap between twin-based and SNP-based heritability is largest at HS+ and narrows thereafter. Plausible contributors to this gap include genetic variation not well tagged by common SNPs, as well as standard twin-model assumptions that may be imperfectly met^29–31^. Our study was not designed to distinguish between these different possibilities. Therefore, rather than making claims about specific biological causes, we treat the gap as a measure of the remaining genetic variance not captured by current common-variant methods for a milestone that nearly everyone achieves.

Several limitations should be considered when interpreting our findings. Residual right-censoring for advanced degrees is a potential issue, particularly in the younger NTR twin sample (mean age 40.9 years), though it is less of a concern in the more mature MoBa cohort (mean age 51.4 years). The external comparison with the EA4 meta-analyses inherits its credential binning and sample composition. As previously discussed, the twin estimates rest on standard assumptions that may be violated, potentially in ways that differ by milestone. Furthermore, generalizability is limited, as our findings reflect dynamics within the specific context of the Norwegian educational system and a primarily European-ancestry population.

The practical implications of our findings concern the appropriate use and interpretation of PGIs for education. A years-based PGI is not uniformly predictive across the educational ladder. Instead, its utility likely peaks for intermediate transitions, such as the entry into higher education, and is diminished at the distributional endpoints: the near-universal high-school transition and the low-prevalence postgraduate thresholds. Our results suggest the appropriate use case is for population-level description and hypothesis testing, not for individual-level differentiation, particularly at these endpoints. Accurate interpretation of PGI-based findings, therefore, requires acknowledging this context-dependent and non-linear performance.

Future research could extend our findings in several specific and feasible ways. First, scale up our single-cohort approach by building milestone-preserving meta-analyses that avoid credential collapse and combine data from multiple cohorts. This would enable the construction of milestone-specific PGIs for out-of-sample evaluation. Second, expand correlation targets beyond EduYears and GCA to include non-cognitive dimensions implicated in educational progression^32^, using designs that limit sample overlap and control for ascertainment. Third, increase resolution by field of study, where genetic correlations and PGI performance may differ in informative ways^33^. Fourth, extend these analyses to multi-ancestry cohorts to test the generalizability of these patterns beyond European-ancestry populations. Fifth, use whole-genome sequencing data to probe the sources of the HS+ twin-SNP gap and to test whether rare or structural variation contributes differentially across milestones. Sixth, incorporate socioeconomic and institutional context explicitly. Compare heritability metrics across tuition-free vs tuition-based systems, changes in financial aid or admissions selectivity, and across parental SES strata. Each step addresses a concrete limitation identified here and will sharpen both interpretation and utility.

Treating education as a series of discrete milestones is essential for accurately mapping its dynamic genetic architecture. This milestone-aware design reveals a different picture at the distributional endpoints: at the near-universal high-school milestone, the twin–SNP gap indicates heritable variance beyond what is captured by common variants alone, while at low-prevalence postgraduate milestones, genetic influences are less aligned with those driving general attainment and cognition, with internal genetic correlations weakening with educational distance. A practical implication is that years-based PGIs and SNP-based heritability are most informative for intermediate attainment and should be applied at the endpoints with caution. Future work that preserves credential granularity, broadens target constructs, and compares findings across diverse educational contexts will yield more accurate inferences about how genetic and environmental influences combine to shape educational trajectories.

## Methods

### Data samples

#### The Norwegian Mother, Father and Child Cohort Study (MoBa)

Our primary discovery analyses were conducted in the Norwegian Mother, Father and Child Cohort Study (MoBa), a population-based pregnancy cohort study conducted by the Norwegian Institute of Public Health^34,35^. Participants were recruited nationwide from 1999-2008 from 50 of Norway’s 52 hospitals with maternity units (41% participation rate), comprising approximately 114,500 children, 95,200 mothers, and 75,200 fathers. Participation required the ability to read Norwegian, as all materials were provided only in Norwegian.

This study analyzed 120,527 participants with registry-linked educational data: 48,953 males and 71,574 females born between 1937 and 1992 (males: 1937-1989; females: 1954-1992; mean birth year 1972.29±5.63 for males, 1974.42±5.08 for females). The mean age of the sample at data extraction (January 2025) was 51.4±5.4 years.

Genotyping was conducted through multiple research projects across 26 batches using various arrays and genotyping centers. Quality control (QC) and imputation followed the MoBaPsychGen pipeline^36^, which included SNP and individual-level QC, accounting for the complex family structure in MoBa. After post-imputation QC, genetic data were available for 207,569 unique individuals (90% of genotyped samples) with 6,981,748 autosomal SNPs. The cohort includes extensive family relationships: 287 monozygotic twin pairs, 22,884 full siblings, 117,004 parent-offspring pairs, and thousands of second- and third-degree relative pairs.

Educational attainment (EA) distribution was: 9.2% (N = 11,129) compulsory education (Comp), 30.0% (N = 36,210) high school (HS), 42.1% (N = 50,724) bachelor’s (BSc), 16.7% (N = 20,187) master’s (MSc), and 1.9% (N = 2,277) doctorate (PhD). Sex-specific distributions showed males with higher Comp (M: 10.9%, F: 8.1%) and HS (M: 38.5%, F: 24.3%) completion, while females achieved higher BSc rates (M: 31.5%, F: 49.3%). MSc rates were similar (M: 16.9%, F: 16.6%), with males slightly higher for PhD (M: 2.2%, F: 1.6%).

The full distribution of EA across birth cohorts and sex in the MoBa sample is shown in Extended Data Fig. 1.

#### The Norwegian Twin Registry (NTR)

To estimate twin-based heritability, we used data from the Norwegian Twin Registry (NTR), a consent-based registry maintained by the Division for Health Data and Digitalization at the Norwegian Institute of Public Health. While the full NTR includes twins born 1915-1991 (N=32,664 individuals), our analytical sample was restricted to twins born 1967-2000 who had completed education questionnaires. Zygosity was determined by questionnaire methods inquiring about twin similarity and has been verified in subsamples by genetic markers and DNA analysis^37^.

The analytical sample included 8,910 twins: 3,054 males (34.3%) and 5,856 females (65.7%), with mean birth years of 1982.83 (SD=10.41) and 1984.70 (SD=10.38), respectively. At the time of data extraction (January 2025), the mean age was 40.9±10.4 years. The sample contained 5,096 monozygotic (MZ) twins (57.2%) and 3,814 dizygotic (DZ) twins (42.8%).

EA distribution was: 12.7% (N = 1,128) Comp, 33.8% (N = 3,012) HS, 37.0% (N = 3,300) BSc, 15.6% (N = 1,391) MSc, and 0.9% (N = 79) PhD. Sex-specific distributions showed similar completion of Comp (M: 12.7%, F: 12.6%), but males completed HS more frequently (M: 38.5%, F: 31.4%), while females earned BSc more often (M: 30.2%, F: 40.6%). Males attained higher rates of MSc (M: 17.4%, F: 14.7%) and PhD (M: 1.2%, F: 0.7%).

The full distribution of EA across birth cohorts and sex in the NTR sample is shown in Extended Data Fig. 2.

### Definition of educational milestones and analytical approaches

EA was classified into five primary categories based on the Norwegian Standard Classification of Education (NUS2000^38^), as detailed in Extended Data Table 1. From these categories, we defined four key educational milestones: achieving at least a high school diploma (HS+), a bachelor’s degree (BSc+), a master’s degree (MSc+), and a doctoral degree (PhD).

To analyze the genetic architecture of these milestones, we employed two complementary analytical frameworks, a transition-specific and a cumulative approach, which are precisely defined in Figure 7.

**Figure 7.**
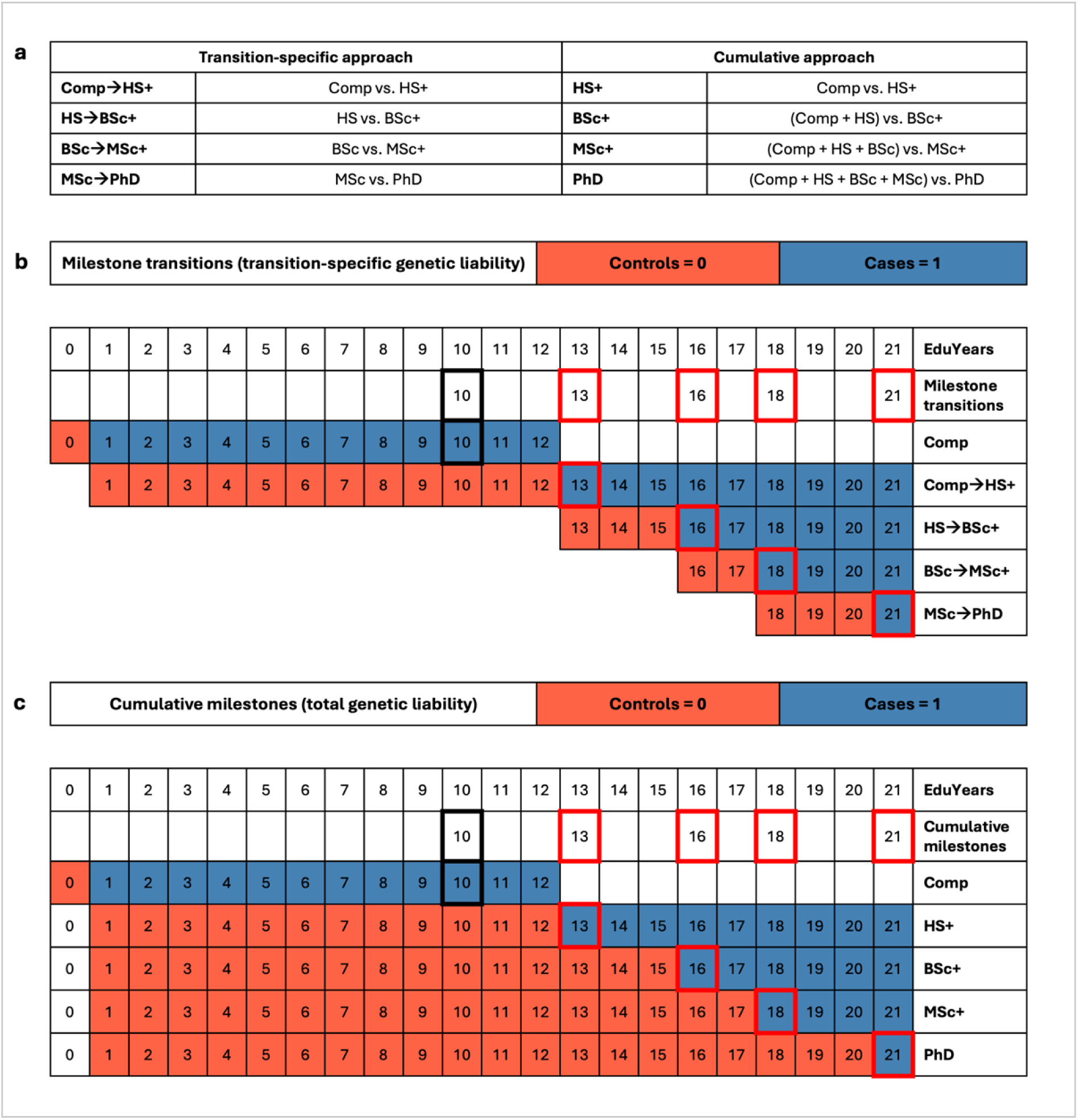
Definitions of transition-specific and cumulative analytical approaches. **a,** A summary of the case-control definitions for each educational milestone under the two primary analytical frameworks used in the study. **b,** Visual illustration of the transition-specific approach, which isolates transition-specific genetic liability. Cases (blue) for each milestone are compared against controls (red) who achieved only the immediately preceding educational level. **c,** Visual illustration of the cumulative approach, which captures the total genetic liability for attaining a milestone. Cases (blue) are compared against all individuals who did not achieve that milestone, regardless of their final attainment.

The transition-specific approach is designed to isolate the genetic factors specific to each educational milestone transition, conditional on having successfully cleared the preceding one. As illustrated in Figure 7b, this framework defines cases as those who advance to a new milestone, while controls are strictly limited to those who achieved only the immediately prior level. For example, to analyze the transition to a bachelor’s degree (HS→BSc+), cases would have a BSc or higher, while controls would have completed high school (HS) but not a BSc. In contrast, the cumulative approach captures the total genetic liability required to reach a given educational milestone. This framework, shown in Figure 7c, compares all individuals who attained a specific milestone or higher (cases) against everyone in the sample who did not reach that level (controls). For instance, in the cumulative analysis of a bachelor’s degree (BSc+), the control group would include all individuals with either compulsory education (Comp) or a high school diploma (HS). The specific sample sizes and effective sample sizes (*N*_eff_) for each analysis are detailed in Table 1.

### Statistical analyses

#### Overview

Our analytical strategy leverages two Norwegian cohorts to characterize genetic influences on educational milestones and is built around a dual framework. We pursued complementary analyses throughout to validate findings through methodological triangulation.

First, under a transition-specific framework, we sought to isolate the genetic factors relevant to each sequential step. To establish empirical ceilings for genetic overlap, we began by conducting an EduYears GWAS in MoBa and correlating it with the EA4 and IQ3 meta-analyses. With these benchmarks established, we conducted our transition-specific milestone GWASs. We used these results to estimate SNP-based heritability (h^2^_SNP_), calculate their genetic correlation (r_g_) with EA4 and IQ3 to identify which transitions diverge from general attainment, and test the predictive power of an EA4-derived polygenic index (PGI).

Second, under a cumulative framework, we conducted a different set of milestone GWASs to model the total genetic liability required to achieve each status. This approach enabled us to assess the genetic overlap (r_g_) between the milestones themselves, using bivariate BOLT-REML to properly handle the complete sample overlap. We also estimated common-variant heritability (h^2^_SNP_) using two converging methods (summary-level LD score regression (LDSC) and individual-level BOLT-REML) to ensure our findings represent biological signal rather than methodological artifacts. Finally, this framework allowed us to contrast our SNP-based results with total additive genetic heritability (h^2^_twin_) from classical twin modeling in the NTR, revealing where additional genetic complexity beyond common variants might exist.

This comprehensive approach allows us to disentangle the genetics of the process of educational progression from the genetics of the final outcome achieved.

#### Genome-wide association studies (MoBa)

We conducted genome-wide association studies (GWAS) in MoBa for EduYears and four educational milestones, with the milestones analyzed using both transition-specific and cumulative definitions (see figure 7 for illustration of definitions and table 1 for sample sizes). We also performed additional exploratory GWAS using non-cumulative contrasts, which directly compared individuals whose highest attainment was a BSc, MSc, or PhD against those whose highest credential was a high school diploma. The EduYears GWAS provided essential benchmarks for establishing reference r_g_ estimates with external meta-analyses, as described in our Statistical Analyses overview.

All GWAS employed identical QC procedures and model specifications. Analyses used fastGWA (GCTA)^40^, which implements a linear mixed model (LMM) with a sparse genetic relationship matrix (GRM) to account for relatedness^41^; population structure was controlled by PCs included as covariates. Although fastGWA was developed for quantitative traits, it provides valid estimates for binary outcomes when population stratification is properly controlled.

The model specification was:

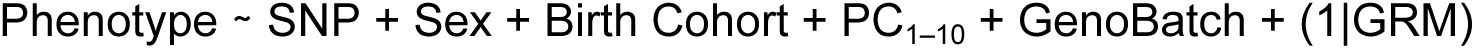

where phenotypes were either continuous (EduYears) or binary (milestone completion). Standard QC excluded SNPs with call rate <0.95, MAF <0.01, imputation INFO <0.80, or Hardy–Weinberg equilibrium p <1×10⁻⁶. Genome-wide significance was set at two-sided p<5×10⁻⁸ ^42^.

Post-GWAS annotation used FUMA^43^ with 250kb window clumping (r²=0.6 for lead SNPs, r²=0.1 for candidate SNPs) using the UK Biobank 10k European panel as the LD reference. Calibration was assessed with LD score regression (LDSC) on HapMap3 SNPs using 1000G-EUR LD scores. Q-Q and Manhattan plots for all analyses are shown in Supplementary Figs. S1–S24. Numerical values for observed-scale SNP-heritability (LDSC) and QC metrics (λGC, LDSC intercept, and attenuation ratio) for each GWAS are provided in Supplementary Table S1. A complete list of genome-wide significant lead variants and their clumped genomic loci is provided in Supplementary Table S2.

Lastly, we used the resulting .fastgwa summary statistics in cross-trait LDSC for external GWAS comparisons, while bivariate BOLT-REML used individual-level data for within-cohort milestone correlations (see subsequent section).

#### Genetic correlations

We estimated pairwise r_g_ to quantify shared genetic influences within cohort, among educational milestones, and between cohort, relating milestones to external GWAS of EA and GCA.

For pairwise within-MoBa r_g_, we fit bivariate BOLT-REML^44^ to cumulative milestone phenotypes. This choice is driven by model identifiability. Transition outcomes are defined on different at-risk subsets (e.g., the PhD transition is defined only among MSc+). In the intersection of two transition risk sets required for bivariate GREML, the earlier transition is constant (always 1) and therefore has zero variance, which places the variance components on the boundary and produces non-convergence. Cumulative outcomes are defined for the full cohort, so both phenotypes retain non-zero variance and the model is identifiable. Consistent with this rationale, effective sample sizes (see Table 1) differ only modestly between cumulative and transition definitions, indicating that the transition non-convergence is not explained by lack of power.

For r_g_ with external GWAS, we used cross-trait LD score regression^45^ to correlate MoBa GWAS, both transition-specific and cumulative definitions, with EA4^23^ and IQ3^25^. We first benchmarked MoBa EduYears against EA4 and IQ3, then estimated r_g_ between each milestone and each external trait. LDSC separates polygenic overlap from confounding using genome-wide LD patterns.

Bivariate BOLT-REML is appropriate for overlapping phenotypes measured on the same individuals when both vary in the joint sample (cumulative outcomes). Cross-trait LDSC is appropriate for between-cohort comparisons from summary statistics (including transition outcomes). Together they provide complementary estimates of genetic sharing across milestones and their links to broader constructs of education and cognition.

#### SNP-based heritability (MoBa)

We estimated common-variant heritability (h²_SNP_) in MoBa using two complementary estimators. (i) BOLT-REML (individual-level): We fit LMMs to QC’d autosomal SNPs (minor-allele frequency > 0.01, imputation INFO > 0.80, Hardy–Weinberg *p*<10^−6^, call rate > 0.95), adjusting for sex, birth-cohort bins, genotyping batch, and PC1–PC10, with a GRM to model relatedness^44^. (ii) LDSC (summary-level): We applied LD Score Regression to fastGWA summary statistics^40^ using 1000G EUR HapMap3 LD scores and report LDSC intercepts/attenuation ratios as calibration checks^26^.

Because outcome definitions differ in identifiability and scale, cumulative and transition phenotypes were handled separately. For cumulative milestones (HS+, BSc+, MSc+, PhD), we report h²_SNP_ from both estimators: LDSC on the cumulative-phenotype GWAS and the diagonal heritability terms from bivariate BOLT-REML. Both procedures return observed-scale heritabilities, which we converted to the liability scale to enable direct comparison with twin-based heritability (h²_twin_). The conversion follows the classical liability-threshold model^46,47^ using the mapping of Lee et al.^27^ as implemented in bigsnpr^48^. We denote the MoBa sample case fraction by *P* and the unconditional population prevalence by *K*; *P* and *K* values for each milestone are reported in Table 1 (MoBa rows for *P*; NTR rows for *K*).

For transition-specific milestones (Comp→HS+, HS→BSc+, BSc→MSc+, MSc→PhD), we estimated h²_SNP_ with LDSC and retain these values on the observed scale. Individual-level GREML was not identifiable because, in the joint analysis set required for a bivariate model, the earlier transition is constant among those at risk, yielding boundary variance components and non-convergence (see Genetic Correlations). Moreover, a liability-scale transform at transitions would produce conditional quantities specific to the at-risk subpopulation and is not directly comparable to the population-level liability estimates reported for cumulative milestones.

To contextualize precision for binary traits, we report effective sample size *N*_eff_ in Table 1, computed as *N*_eff_ = 4/(1/*N*_cases_+1/*N*_controls_) = 4*NP*(1−*P*), a standard proxy for power^39^. *N*_eff_ is used for interpretation only and does not enter the liability-scale conversion.

Using both BOLT-REML and LDSC for cumulative outcomes provides triangulation: REML is statistically efficient but sensitive to covariate control for structure, whereas LDSC is robust to such confounding via the intercept, albeit typically less efficient. Agreement between the two therefore increases confidence that the reported values reflect common-variant contributions to the cumulative milestones.

#### Polygenic indices (MoBa)

In MoBa, we constructed genome-wide PGIs PRS-CS^49^ using summary statistics from EA4, excluding the 23andMe cohort because their data are not publicly available^23^. PRS-CS applies a Bayesian continuous-shrinkage prior and accounts for linkage disequilibrium through an external LD reference panel, which permits genome-wide weighting without *p*-value thresholding. After allele alignment and standard QC in the target data, individual PGI scores were computed and standardized to mean 0 and variance 1 within MoBa.

We evaluated the relation between the EA4 PGI and educational milestones in two complementary ways and under both transition-specific and cumulative definitions. First, we compared mean PGI levels between those who advanced and those who did not advance to each milestone transition; this assesses genetic stratification across the schooling sequence. Second, we quantified predictive accuracy in multivariable probit models that regressed each binary milestone on the standardized PGI, adjusting for sex, birth-cohort bins, and the first ten PCs.

For predictive accuracy we report two metrics that intentionally live on different, interpretable scales. Tjur’s coefficient of discrimination^50^ is reported on the observed scale as the difference in mean predicted probabilities between cases and controls; uncertainty is summarized with nonparametric bootstrap 95% confidence intervals obtained by refitting the models on resampled data (1000 iterations).

For cumulative outcomes only, we also report a liability-scale PGI metric that captures the variance of liability explained by the PGI, obtained from the probit slope using a standardized PGI that was first residualized on the same covariates used in the prediction model; this follows the mapping described by Lee et al.^27^, also with bootstrapped (1,000 resamples) 95% confidence intervals.

We do not convert Tjur’s R^2^ to the liability scale because such a transformation is not standard and is not directly comparable to the liability-variance metric. For transition-specific outcomes, we keep PGI metrics on the observed scale and pair them with observed-scale SNP-based heritability estimates to maintain internal consistency within the conditional framework.

Together, the mean-difference analyses and the two prediction metrics address related but distinct questions: the former captures between-group stratification by milestone, while the latter assesses individual-level discrimination. We interpret these PGI results alongside SNP-based and twin-based heritability estimates to contextualize the explanatory power of the EA-based PGIs.

#### Twin-based heritability (NTR)

We analyzed the NTR sample using a classical twin design^51,52^ within a liability-threshold framework, which is appropriate for the binary milestone outcomes. This approach decomposes the variance in liability into additive genetic (A), shared environmental (C), and unique environmental (E) components (the ACE model).

We implemented these models in Stan^53^, a platform for Bayesian statistical modeling, to obtain posterior distributions for the ACE parameters, from which we derived point estimates and 95% credible intervals. This Bayesian approach provides more robust uncertainty quantification than traditional maximum likelihood methods, particularly important given the varying prevalences of our educational milestones.

Due to insufficient statistical power and potential model instability for analyzing specific transitions within the NTR sample, our twin analyses were restricted to the cumulative definition of educational milestones. The transition-specific approach would require analyzing twins discordant for consecutive educational levels (e.g., one twin stopping at high school while the other continues to bachelor’s), which yields very small cell sizes given the correlation of EA within twin pairs.

The resulting additive genetic heritability (h²_twin_) and the C and E components are all reported on the unobserved liability scale. These estimates theoretically capture all additive genetic variation, including contributions from rare variants, structural variants, and gene-gene interactions that are invisible to SNP-based methods. Comparing h²_twin_ with h²_SNP_ thus reveals the so-called missing heritability at each educational milestone. Full details on the model specification, choice of priors, and MCMC settings are provided in Supplementary Note 2.

## Data Availability

All genome-wide association study (GWAS) summary statistics generated in this study will be deposited in the NHGRI-EBI GWAS Catalog (accession numbers pending) upon publication. The summary statistics used for external comparisons, EA4 and IQ3, are available for download from the Social Science Genetic Association Consortium (SSGAC) upon user registration, as detailed in their original publications.

The individual-level genetic and phenotypic data for this study were obtained from the Norwegian Mother, Father and Child Cohort Study (MoBa) and the Norwegian Twin Registry (NTR). Due to Norwegian privacy laws and the sensitive nature of the data, these resources are under restricted access and cannot be shared directly by the authors. Access to these datasets for replication or other research purposes can be obtained by qualified researchers through a formal application process to the respective data holders, subject to ethical review and data protection regulations. Information on applying for access is available from the Norwegian Institute of Public Health.

## Code Availability

The full Stan code for implementing the Bayesian hierarchical twin model is provided in Supplementary Note 2. Scripts used for genome-wide association studies (GWAS) and polygenic index (PGI) analyses relied on publicly available software, with key commands and parameters detailed in the Methods section. While the R scripts themselves are not publicly posted due to their integration with restricted data, the open-source nature of the software cited ensures that the analytical procedures are transparent and replicable.

## Acknowledgements

We are grateful to all the participating families in Norway who take part in this on-going cohort study. We also thank the twins in the Norwegian Twin Registry (NTR) for their invaluable contributions. Educational register data were provided by Statistics Norway (SSB). The interpretations and conclusions presented here are those of the authors and do not necessarily reflect the views of NIPH, SSB, or the funding agencies.

The MoBa, NTR and SSB data were used via the project SUBPU, which is approved by the Regional Committees for Medical and Health Research Ethics (ref. 2017/2205). The University of Oslo is responsible for the data handling in SUBPU and has conducted a Data Protection Impact Assessment (DPIA) in collaboration with the Norwegian Agency for Shared Services in Education and Research (Sikt; ref. 962088). The data access and management of SUBPU is financed by the Research Council of Norway (RCN), the European Research Council (ERC), and the Department of Psychology at the University of Oslo. (Data were provided as pseudonymized research extracts by the data controllers and analysed within UiO’s secure environment; only aggregate, non-identifiable results left the secure environment.)

This work was performed on the TSD (Tjenester for Sensitive Data) facilities, owned by the University of Oslo, operated and developed by the TSD service group at the University of Oslo, IT-Department (USIT). Computations were performed on resources provided by Sigma2 – the National Infrastructure for High-Performance Computing and Data Storage in Norway (ref. NS9867S).

For generating high-quality genomic data, we thank the Norwegian Institute of Public Health (NIPH), the HARVEST collaboration, the NORMENT Centre at the University of Oslo, the Center for Diabetes Research at the University of Bergen, deCODE Genetics, the Research Council of Norway, the South-Eastern and Western Norway Regional Health Authorities, the ERC AdG, Stiftelsen KG Jebsen, the Trond Mohn Foundation, and the Novo Nordisk Foundation.

This work was supported by the Research Council of Norway (RCN #336078, #288083, #314601), the European Research Council (ERC) (#101045526, #818425, #101088481, #818420), and the Department of Psychology, University of Oslo. It was also supported by UiO:Life Science through the AHeadForLife convergence environment (2022–2026), and by the Research Council of Norway through the AHeadForLife project (grant no. 325001) to K.B.W. We thank Perline A. Demange for helpful comments on an early version of the manuscript presented at the UiO midterm evaluation.

## Author contributions

**E.H.K.:** Conceptualization, Methodology, Formal Analysis, Software, Data Curation, Visualization, Writing – Original Draft.

**Y.W.:** Methodology, Formal Analysis, Software, Funding Acquisition, Supervision, Writing – Review & Editing.

**K.B.W.:** Funding Acquisition, Resources, Writing – Review & Editing.

**T.H.L.:** Conceptualization, Software, Funding Acquisition, Resources, Writing – Review & Editing.

**O.R.:** Conceptualization, Methodology, Formal Analysis, Software, Funding Acquisition, Supervision, Resources, Writing – Review & Editing.

## Footnotes

* Examples include the American Community Survey ^8^ and the European Social Survey ^9^.

