## Supplementary_Information for "Beyond years of schooling: Shifting genetic influences across educational milestones in two Norwegian cohorts"

**Supplementary figures (S1-S26) 2**
Transition-specific GWAS results and QC (S1-S8) 2
S1: Manhattan plot, milestone transition Comp→HS+ 2
S2: QQ plot, milestone transition Comp→HS+ 2
S3: Manhattan plot, milestone transition HS→BSc+ 3
S4: QQ plot, milestone transition HS→BSc+ 3
S5: Manhattan plot, milestone transition BSc→MSc+ 4
S6: QQ plot, milestone transition BSc→MSc+ 4
S7: Manhattan plot, milestone transition MSc→PhD 5
S8: QQ plot, milestone transition MSc→PhD 5
Cumulative GWAS results and QC (S9-S16) 6
S9: Manhattan plot, cumulative milestone HS+ 6
S10: QQ plot, cumulative milestone HS+ 6
S11: Manhattan plot, cumulative milestone BSc+ 7
S12: QQ plot, cumulative milestone BSc+ 7
S13: Manhattan plot, cumulative milestone MSc+ 8
S14: QQ plot, cumulative milestone MSc+ 8
S15: Manhattan plot, cumulative milestone PhD 9
S16: QQ plot, cumulative milestone PhD 9
Contrast GWAS results and QC (S17-S22) 10
S17: Manhattan plot, contrast (non-cumulative) BSc vs. HS 10
S18: QQ plot, contrast (non-cumulative) BSc vs. HS 10
S19: Manhattan plot, contrast (non-cumulative) MSc vs. HS 11
S20: QQ plot, contrast (non-cumulative) MSc vs. HS 11
S21: Manhattan plot, contrast (non-cumulative) PhD vs. HS 12
S22: QQ plot, contrast (non-cumulative) PhD vs. HS 12
EduYears GWAS results and QC (S23-S24) 13
S23: Manhattan plot, EduYears 13
S24: QQ plot, EduYears 13
Additional analyses (S25-S26) 14
S25: Genetic correlations between cumulative milestones and EA4/IQ3 14
S26: ACE decomposition for cumulative milestones 14
**Supplementary Notes 1-2 15**
Supplementary Note 1: 15
Composition and Measurement Limitations in the EA4 Comparator GWAS 15
Supplementary Note 2: 16
Methodological Details for the Bayesian Twin Model 16
1. Model Specification 16
2. Hierarchical Structure: Sex- and Cohort-Specific Thresholds 16
3. Choice of Priors 17
4. MCMC Settings and Implementation 17
5. Stan Model Code 18

### Supplementary figures (S1-S26)

#### Transition-specific GWAS results and QC (S1-S8)

##### S1: Manhattan plot, milestone transition Comp→HS+

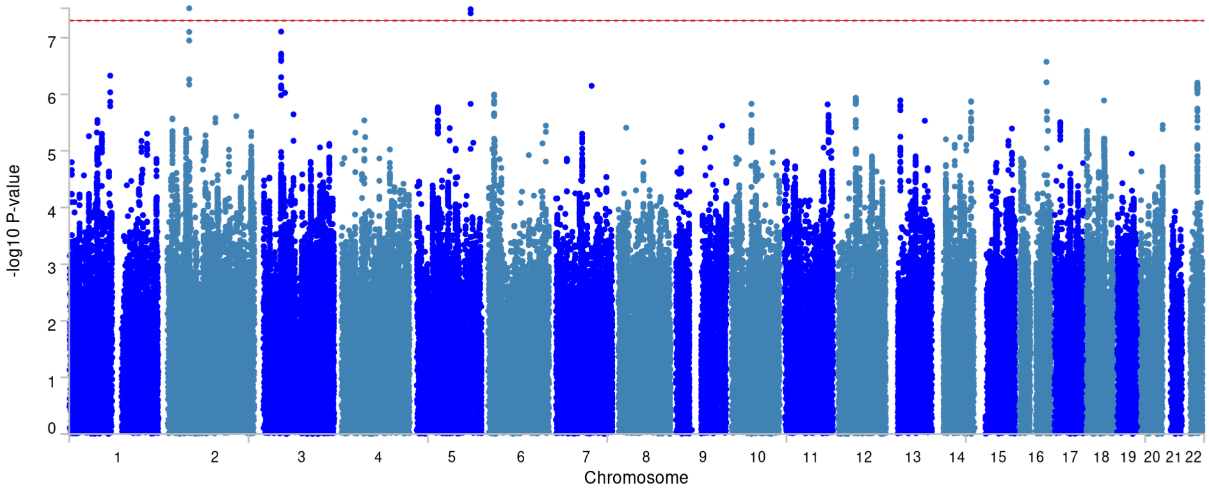


##### S2: QQ plot, milestone transition Comp→HS+

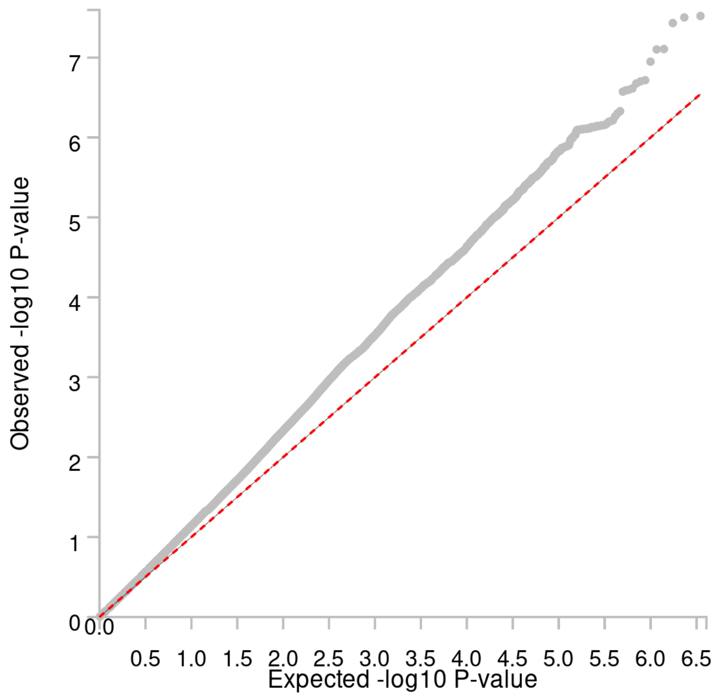

S3: Manhattan plot, milestone transition HS→BSc+

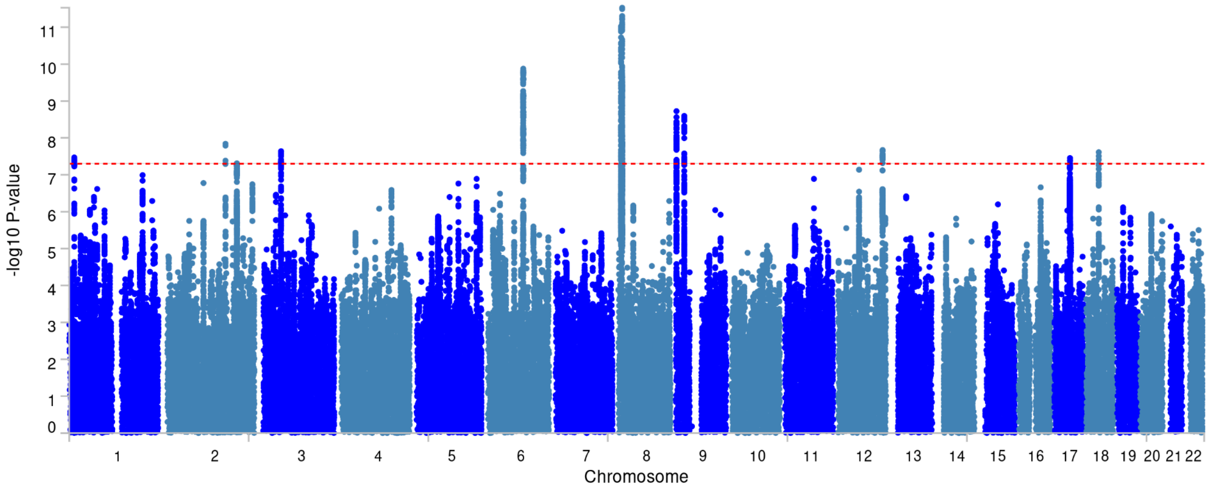


S4: QQ plot, milestone transition HS→BSc+

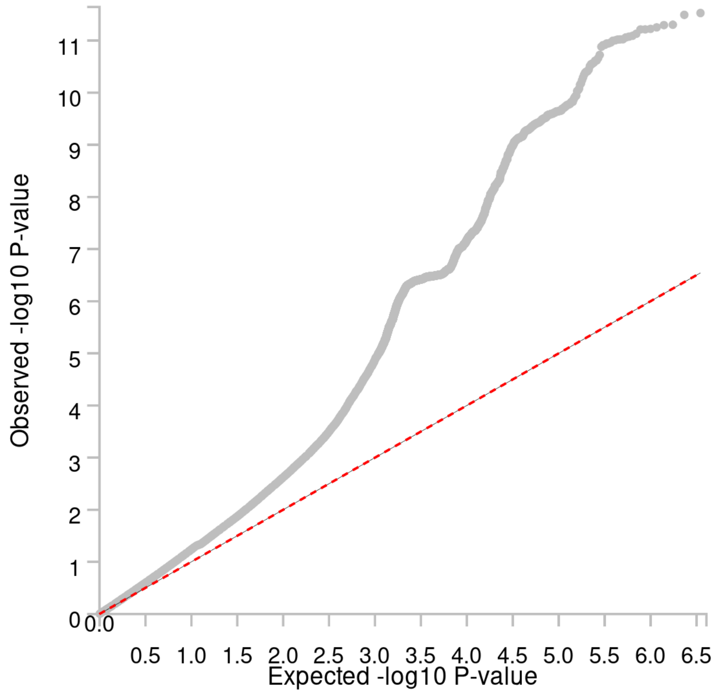

S5: Manhattan plot, milestone transition BSc→MSc+

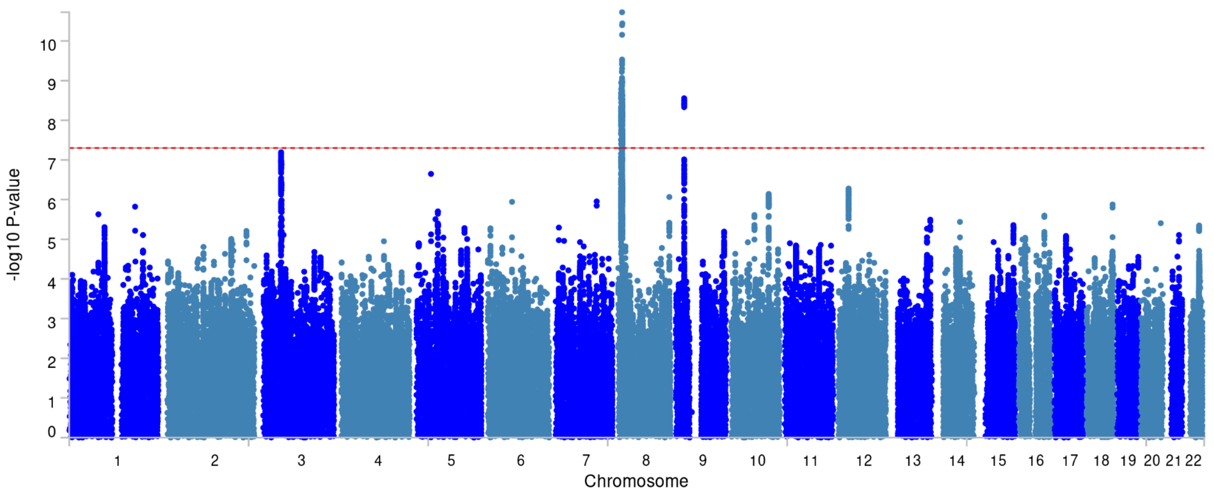


S6: QQ plot, milestone transition BSc→MSc+

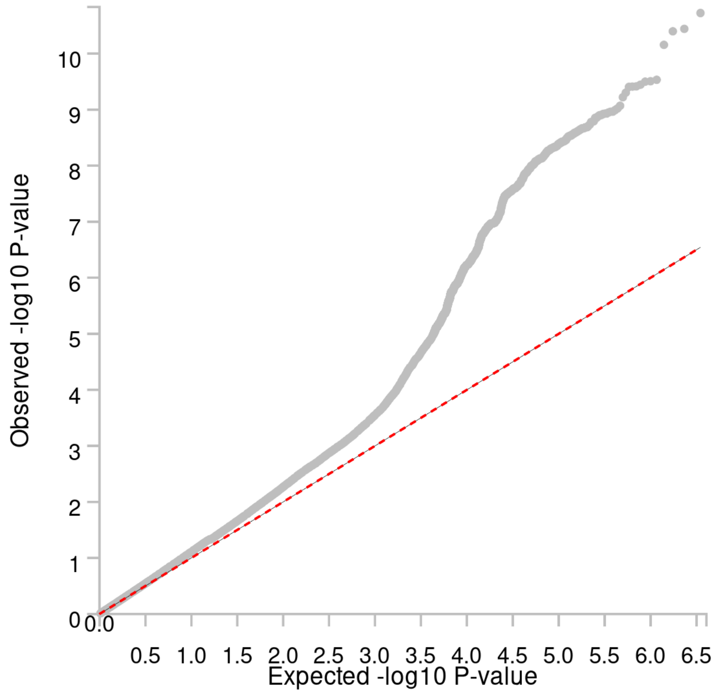

#### S7: Manhattan plot, milestone transition MSc→PhD

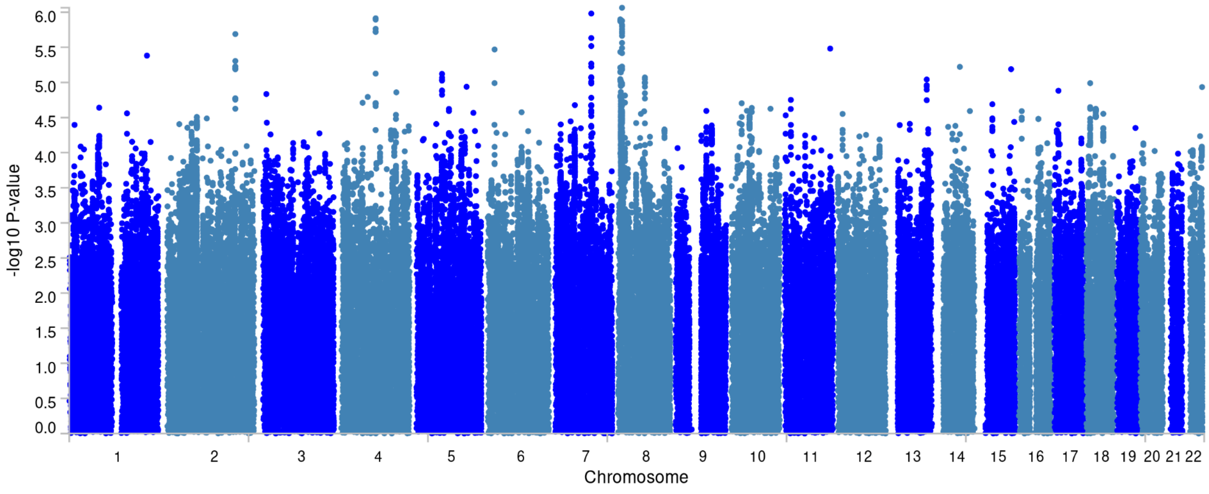


#### S8: QQ plot, milestone transition MSc→PhD

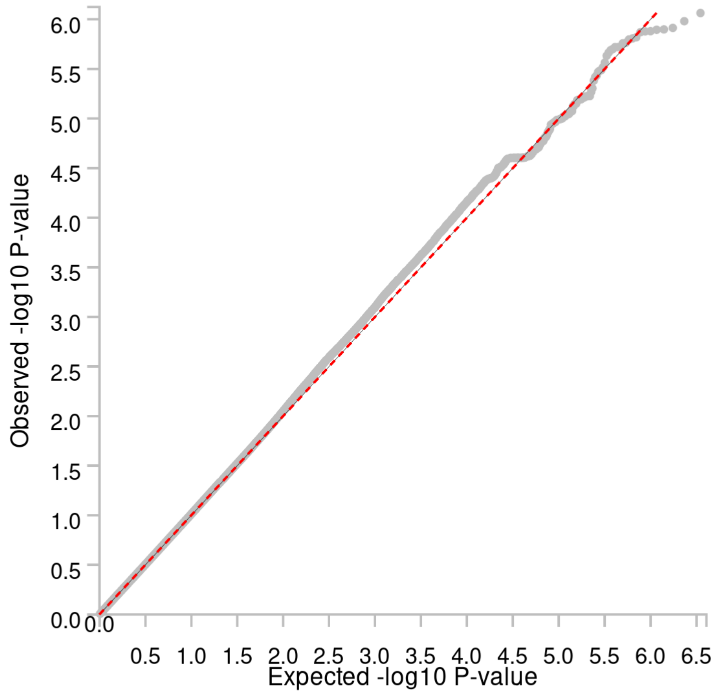

### Cumulative GWAS results and QC (S9-S16)

#### S9: Manhattan plot, cumulative milestone HS+

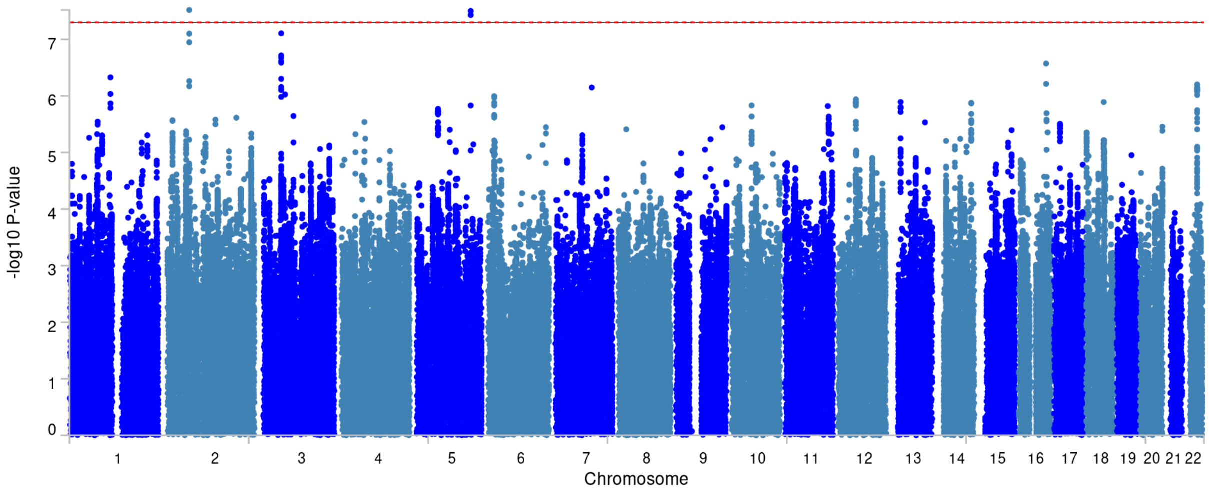


#### S10: QQ plot, cumulative milestone HS+

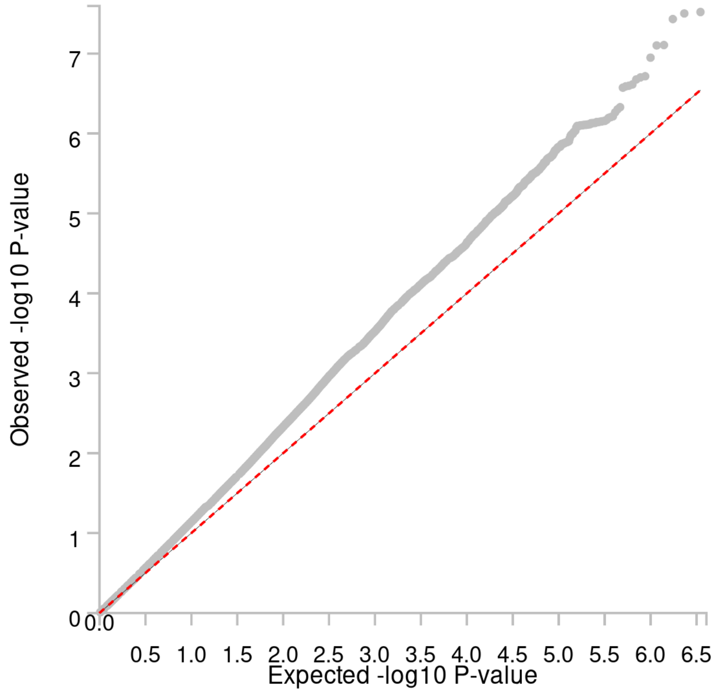

#### S11: Manhattan plot, cumulative milestone BSc+

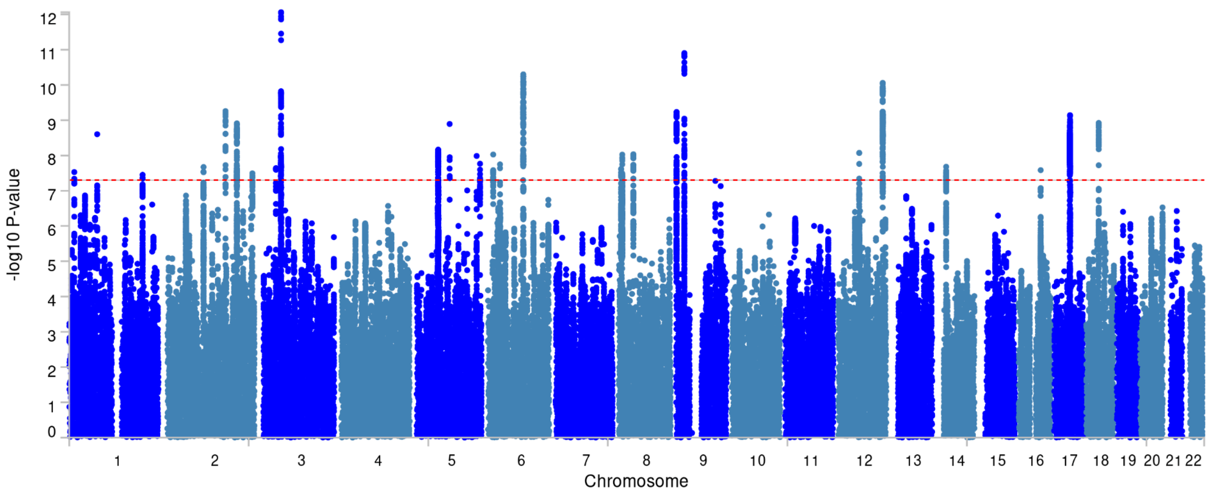


#### S12: QQ plot, cumulative milestone BSc+

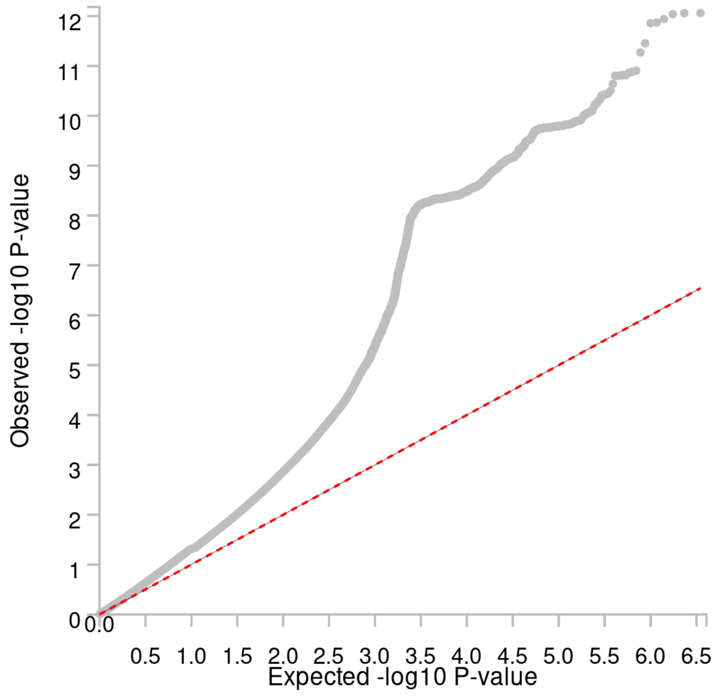

#### S13: Manhattan plot, cumulative milestone MSc+

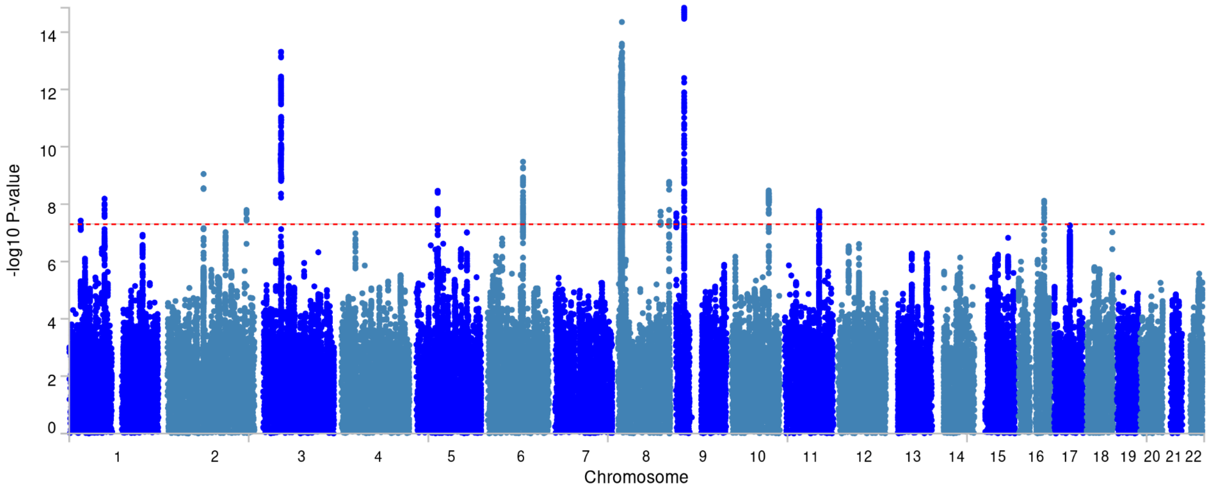


#### S14: QQ plot, cumulative milestone MSc+

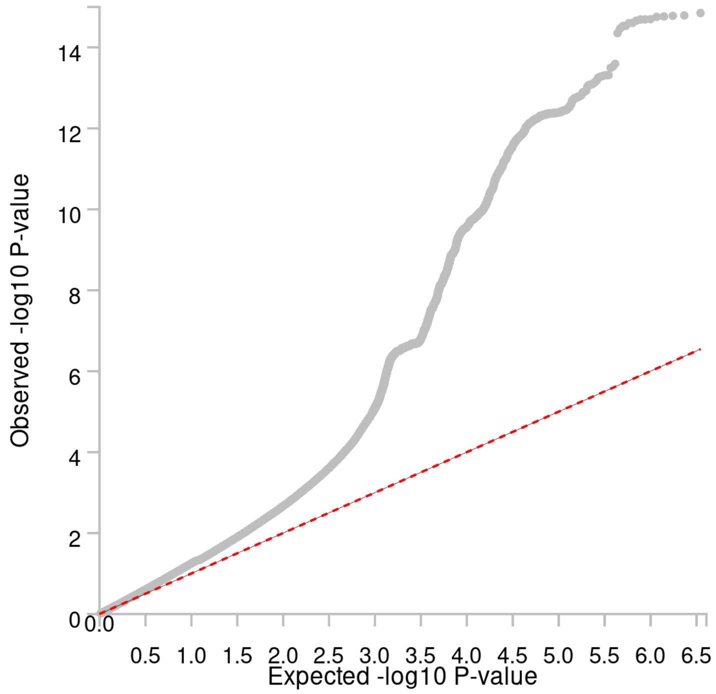

S15: Manhattan plot, cumulative milestone PhD

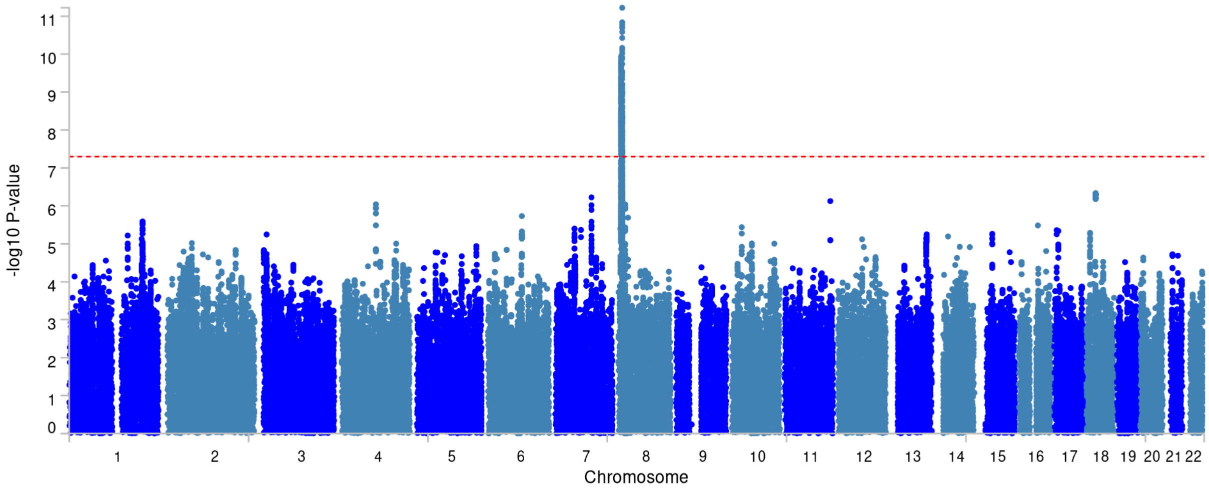


S16: QQ plot, cumulative milestone PhD

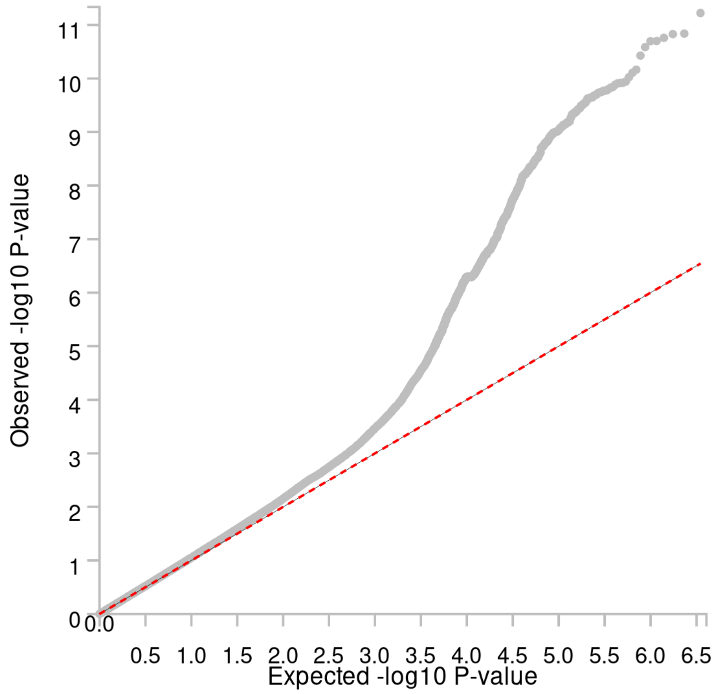

### Contrast GWAS results and QC (S17-S22)

#### S17: Manhattan plot, contrast (non-cumulative) BSc vs. HS

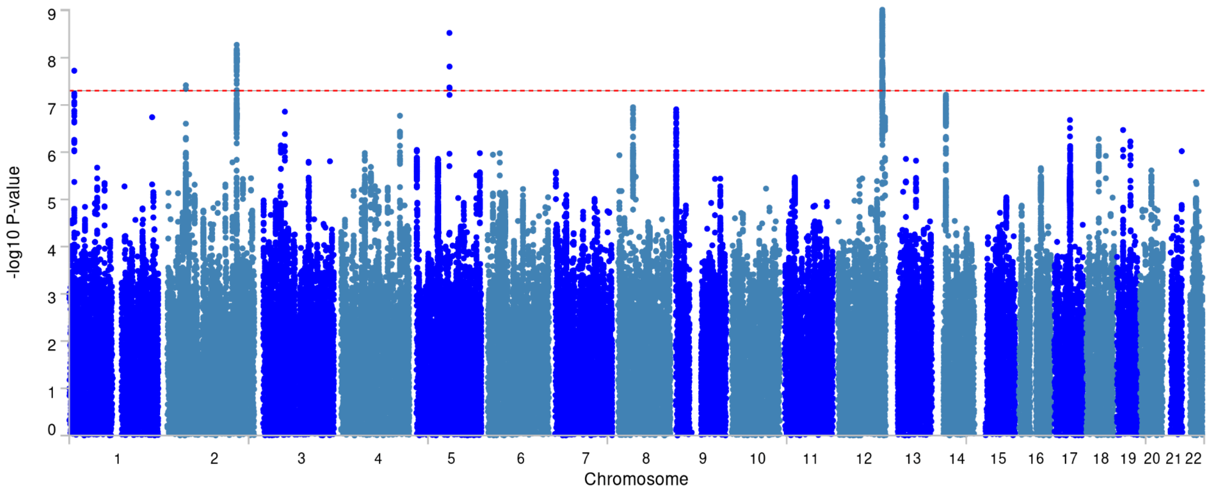


#### S18: QQ plot, contrast (non-cumulative) BSc vs. HS

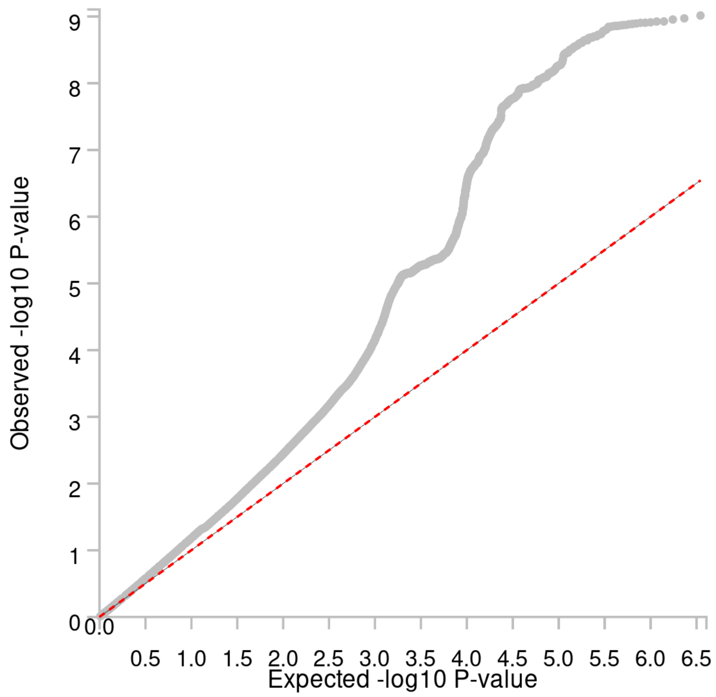

#### S19: Manhattan plot, contrast (non-cumulative) MSc vs. HS

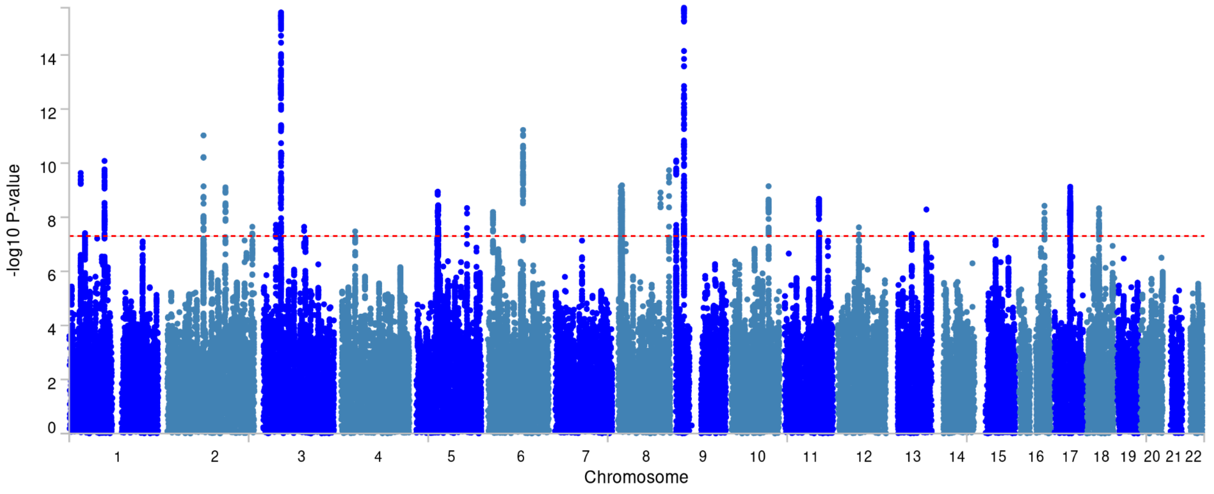


#### S20: QQ plot, contrast (non-cumulative) MSc vs. HS

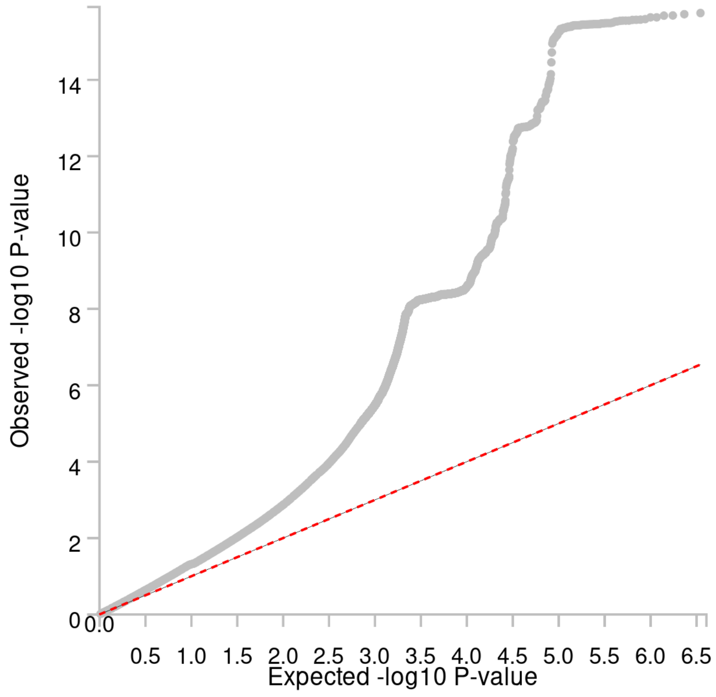

#### S21: Manhattan plot, contrast (non-cumulative) PhD vs. HS

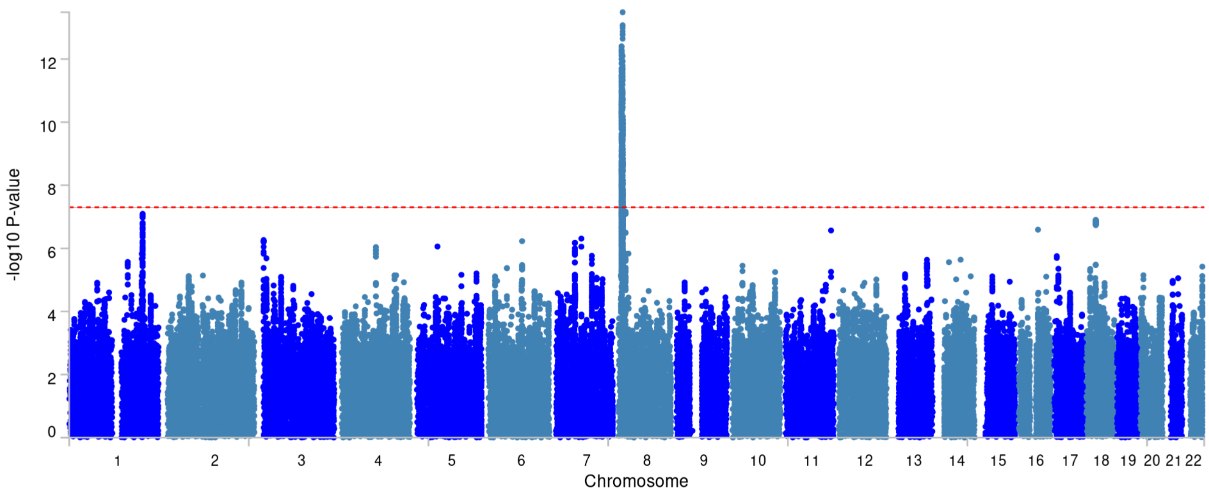


#### S22: QQ plot, contrast (non-cumulative) PhD vs. HS

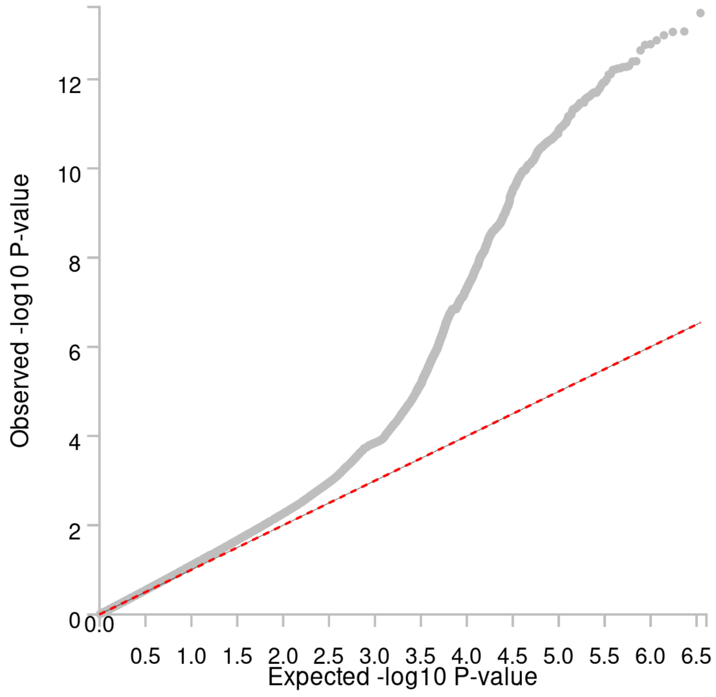

### EduYears GWAS results and QC (S23-S24)

#### S23: Manhattan plot, EduYears

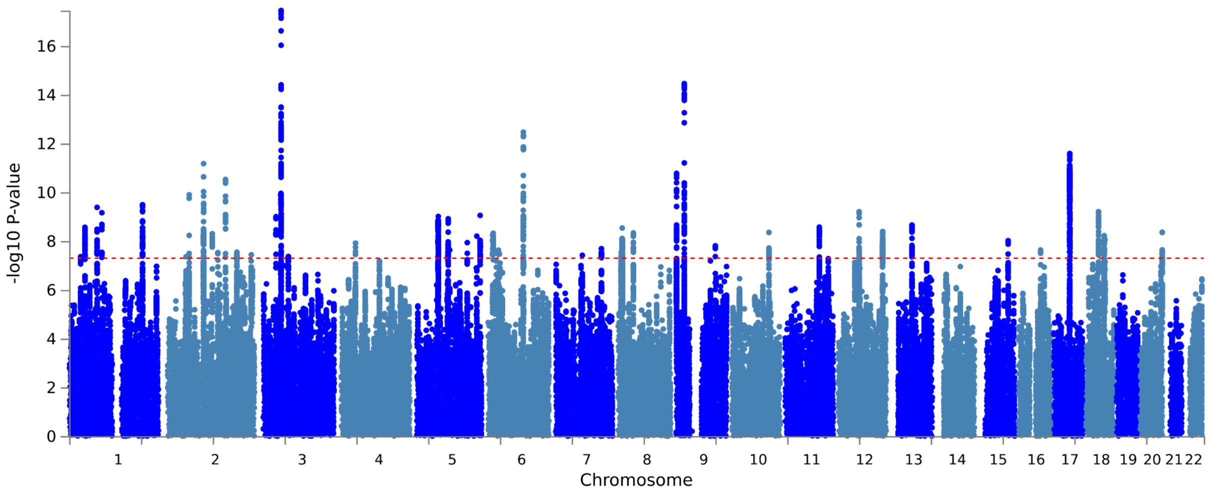


#### S24: QQ plot, EduYears

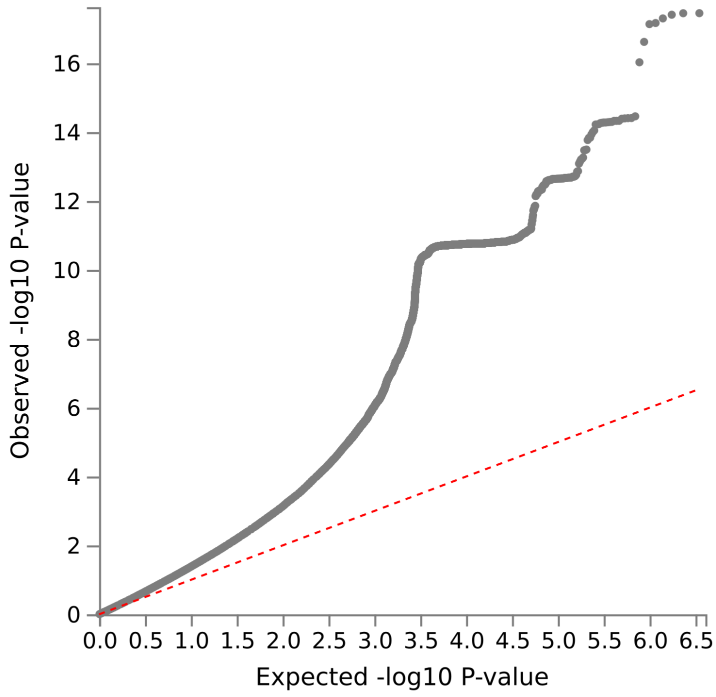

### Additional analyses (S25-S26)

#### S25: Genetic correlations between cumulative milestones and EA4/IQ3

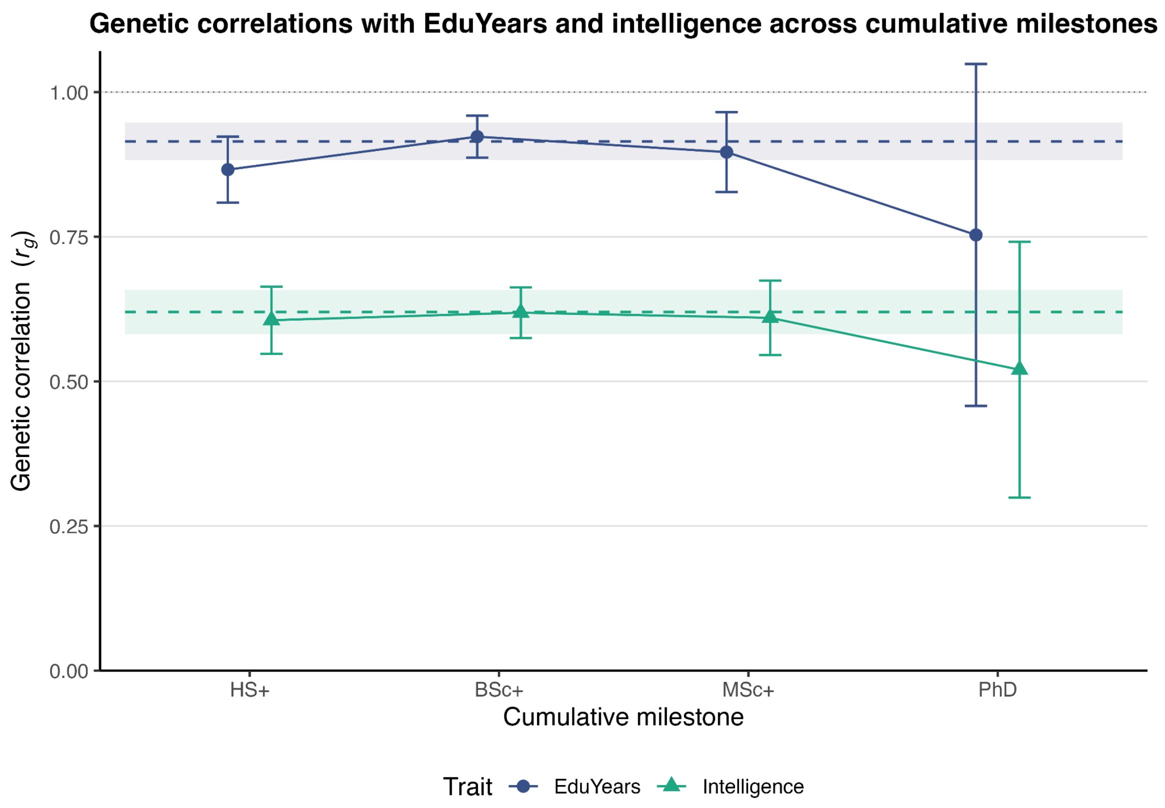


#### S26: ACE decomposition for cumulative milestones

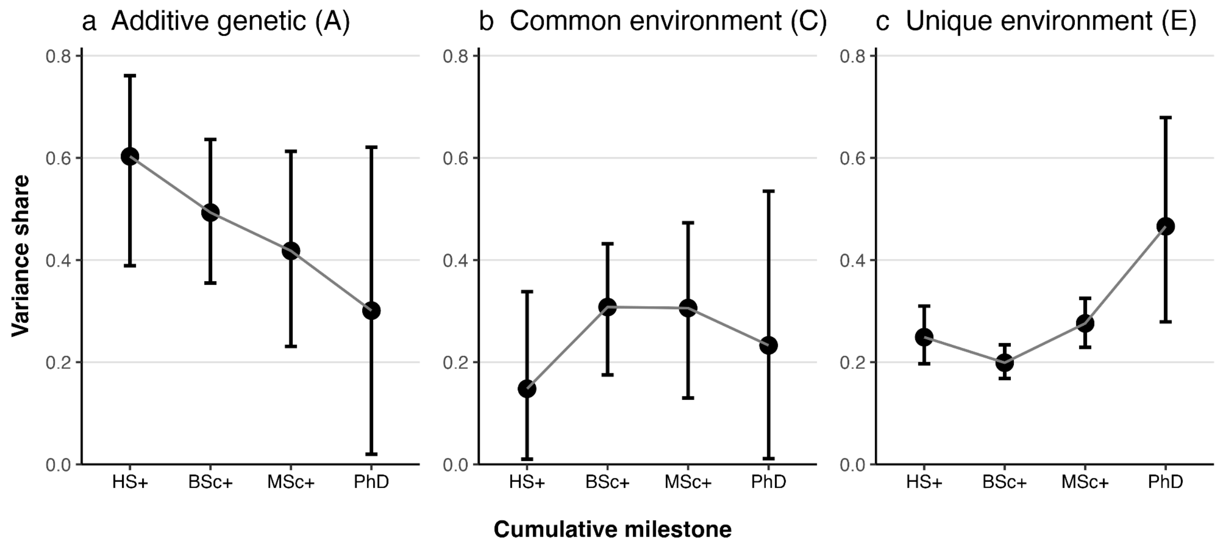

### Supplementary Notes 1-2

#### Supplementary Note 1:

##### Composition and Measurement Limitations in the EA4 Comparator GWAS

Our analyses of external genetic correlations ($r_g$) with educational attainment (EA) and the performance of the EA4-derived polygenic index (PGI) rely on summary statistics from the EA4 meta-analysis. The interpretation of our findings, particularly the attenuated correlations at postgraduate milestones, requires a clear understanding of the composition and measurement properties of the specific EA4 subset available for these comparisons.

Due to data access restrictions, our analyses necessarily excluded summary statistics from the 23andMe cohort, which constitutes approximately 75% of the total EA4 sample size. The remaining, publicly available EA4 subset (N ≈ 765,283) has a distinct composition. The UK Biobank cohort is the largest single contributor, representing approximately 58% of this subset (N ≈ 441,121). Within the UK Biobank, educational attainment is measured with a single binary survey item that distinguishes only whether a participant obtained any college or university degree (Okbay et al., 2022; see their Supplementary Note, Section 1).

The remaining 42% of the subset comprises 69 different cohorts (total N ≈ 324,162) with more varied and often more granular measurement approaches (Lee et al., 2018; see their Supplementary Tables 16, 17, and 22). However, the harmonization of these diverse measures for the EA4 meta-analysis relied on the International Standard Classification of Education (ISCED) 1997 framework (UNESCO, 2006). A key feature of this framework is that it collapses bachelor's and master's degrees into a single category (ISCED level 5), which is then mapped to a single value of 19 years of education in the EduYears metric.

These measurement limitations have direct implications for our findings. Our study, using Norwegian registry data, precisely distinguishes between bachelor's, master's, and doctoral degrees. In contrast, the EA4 comparator GWAS collapses the bachelor's-master's distinction for all participants and uses a simple binary degree measure for the majority of its sample. This structural compression of postgraduate credentials in the EA4 data means that genetic variants specifically associated with completing a master's degree are necessarily pooled with those for a bachelor's degree. This will mechanically dilute the genetic correlation for our BSc→MSc+ transition, as the EA4 summary statistics cannot capture genetic factors unique to this specific step. Similarly, while EA4 does distinguish doctoral degrees, the predominance of the binary degree measure in its largest cohort likely weakens the genetic signal specific to this highest educational transition as well.

### Supplementary Note 2:

#### Methodological Details for the Bayesian Twin Model

This note provides detailed information on the specification, priors, and implementation of the Bayesian hierarchical twin model used to estimate heritability for the four educational milestones.

##### 1. Model Specification

We implemented a liability-threshold model to analyze the binary educational attainment outcomes for same-sex monozygotic (MZ) and dizygotic (DZ) twin pairs. The model assumes that an underlying continuous liability determines the binary outcome (i.e., whether an individual achieves a given educational milestone). An individual *i*'s liability ($L_i$) is decomposed into additive genetic (*A*), shared environmental (*C*), and unique environmental (*E*) components:

$L_i = A_i + C_i + E_i$

The total variance of the liability is standardized to 1, such that:

$Var(A) + Var(C) + Var(E) = h^2 + c^2 + e^2 = 1.$

The parameters $h^2$, $c^2$, and $e^2$ represent the proportions of variance explained by *A*, *C*, and *E*, respectively.

For a given twin pair, the model leverages the genetic relatedness of MZ twins (who share ~100% of their segregating genes) and DZ twins (who share ~50% on average). The shared component of liability for an MZ pair is $\sqrt{h^2 + c^2}$, while for a DZ pair it is $\sqrt{0.5h^2 + c^2}$. The unique (residual) standard deviation for an individual in an MZ pair is $\sqrt{e^2}$, and for an individual in a DZ pair it is $\sqrt{e^2 + 0.5h^2}$ .

##### 2. Hierarchical Structure: Sex- and Cohort-Specific Thresholds

To account for secular trends and gender differences in educational attainment across birth cohorts, we modeled the liability threshold as a hierarchical parameter that varied by both sex and cohort. The model estimates separate thresholds for males and females within each birth cohort.

This was implemented as a random walk, where the threshold for a given sex in a specific cohort is determined by a global mean for that sex plus a cumulative deviation specific to that cohort. This flexible approach allows the prevalence of achieving an educational milestone to change non-linearly over time for males and females independently, without imposing a specific functional form on the trend.

##### 3. Choice of Priors

We used weakly informative priors to regularize the model and ensure stable estimates, while allowing the data to drive the posterior distributions.

- Variance Components (*ACE*): A Dirichlet prior was placed on the simplex of the three variance components:

  *ACE ~ Dirichlet* (1,1,1)

  This is a uniform prior over the space of possible proportions for $h^2$, $c^2$, and $e^2$.

- Global Thresholds (*threshold_global*): The global mean thresholds for males and females were given a standard normal prior:

  *Threshold_global ~ Normal*(0,1)

- Threshold Variation (*threshold_sigma*): The standard deviation of the annual change in thresholds for each sex was given a half-normal prior, weakly centered around zero:

  *Threshold_sigma ~ Normal*(0, 0.2)

- **Standardized Parameters**: All non-centered standardized parameters (*mz_std, dz_std, threshold_std*) were given standard normal priors, *N*(0,1), a standard practice in hierarchical Bayesian models.

##### 4. MCMC Settings and Implementation

The model was written in Stan, a probabilistic programming language, and posterior distributions were estimated using the No-U-Turn Sampler (NUTS), an efficient variant of Hamiltonian Monte Carlo. The analysis was implemented in R using the *rstan* package (version 2.32.6, interfacing with Stan version 2.32.2).

For each of the four educational milestones (HS+, BSc+, MSc+, and PhD), we ran a separate model.

- For all the models, HS+, BSc+, MSc+, and PhD, we ran 8 parallel Markov chains for 6,000 iterations each, with the first 2,000 iterations discarded as warmup. This resulted in 32,000 post-warmup samples for estimating the posterior distributions.

Convergence of the chains was assessed by ensuring the Gelman-Rubin statistic $\widehat{R}$ was below 1.01 for all parameters, observing no divergent transitions after warmup, and by visual inspection of the trace plots to check for good mixing and stationarity.

**5. Stan Model Code**

```
functions {
  real log_prob_bin(int twin_cat, real twinpair_mu, real threshold, real residual_sd){
    real temp_sum = 0;
    if (twin_cat == 0){
      temp_sum = normal_lcdf(threshold | twinpair_mu, residual_sd);
    } else {
      temp_sum = log1m_exp(normal_lcdf(threshold | twinpair_mu, residual_sd));
    }
    return temp_sum;
  }
}

data {
  int<lower=1> mz_twinpairs_n;
  int<lower=1> dz_twinpairs_n;
  int<lower=1> T; // number of unique cohorts
  int<lower=1, upper=T> mz_cohort[mz_twinpairs_n]; // Cohort index for each mz twin pair
  int<lower=1, upper=T> dz_cohort[dz_twinpairs_n]; // Cohort index for each dz twin pair
  int<lower=0, upper=1> mz_y[mz_twinpairs_n, 2]; // Outcome for mz twin pairs
  int<lower=0, upper=1> dz_y[dz_twinpairs_n, 2]; // Outcome for dz twin pairs
  int<lower=0, upper=1> sex_mz[mz_twinpairs_n]; // Sex for mz twin pairs
  int<lower=0, upper=1> sex_dz[dz_twinpairs_n]; // Sex for dz twin pairs
}

parameters {
  // Heritability parameters
  simplex[3] ACE;
  // Shared components
  vector[mz_twinpairs_n] mz_std;
  vector[dz_twinpairs_n] dz_std;
  // Educational thresholds - male and female separately
  vector[2] threshold_global; // Global threshold
  vector<lower = 0>[2] threshold_sigma; // Global threshold - SD of annual change
  matrix[T-1, 2] threshold_std; // cohort specific annual CHANGE in threshold
```

```
}

transformed parameters {
  real mz_shared_sd;
  real dz_shared_sd;
  real mz_unique_sd;
  real dz_unique_sd;
  vector[mz_twinpairs_n] mz_re;
  vector[dz_twinpairs_n] dz_re;
  vector[T] threshold_male;
  vector[T] threshold_female;

  // Calculate the shared and unique standard deviations
  mz_shared_sd = sqrt((ACE[1] + ACE[2]));
  dz_shared_sd = sqrt((0.5 * ACE[1] + ACE[2]));
  mz_unique_sd = sqrt(ACE[3]);
  dz_unique_sd = sqrt(ACE[3] + 0.5 * ACE[1]);

  // Random effects for each twin pair
  mz_re = mz_shared_sd * mz_std;
  dz_re = dz_shared_sd * dz_std;

  // Calculate the thresholds for each cohort and sex
  {
    vector[T] temp_vec = cumulative_sum(append_row(0,threshold_std[,1]));
    threshold_male = threshold_global[1] + threshold_sigma[1] * (temp_vec - mean(temp_vec));

    temp_vec = cumulative_sum(append_row(0,threshold_std[,2]));

    threshold_female = threshold_global[2] + threshold_sigma[2] * (temp_vec - mean(temp_vec));
  }
}

model {
  // Priors
  ACE ~ dirichlet(rep_vector(1, 3));
  mz_std ~ std_normal();
  dz_std ~ std_normal();
  threshold_global ~ normal(0, 1);
  threshold_sigma ~ normal(0, 0.2);
  to_vector(threshold_std) ~ std_normal();

  // Likelihood for MZ twins
  for (n in 1:mz_twinpairs_n) {
    int t = mz_cohort[n];
```

```
    real threshold = (sex_mz[n] == 0) ? threshold_male[t] : threshold_female[t]; // Use sex_mz for MZ
twins
    target += log_prob_bin(mz_y[n, 1], mz_re[n], threshold, mz_unique_sd);
    target += log_prob_bin(mz_y[n, 2], mz_re[n], threshold, mz_unique_sd);
  }

  // Likelihood for DZ twins
  for (n in 1:dz_twinpairs_n) {
    int t = dz_cohort[n];
    real threshold = (sex_dz[n] == 0) ? threshold_male[t] : threshold_female[t]; // Use sex_dz for DZ
twins
    target += log_prob_bin(dz_y[n, 1], dz_re[n], threshold, dz_unique_sd);
    target += log_prob_bin(dz_y[n, 2], dz_re[n], threshold, dz_unique_sd);
  }
}
```
