## Extended_Data for "Beyond years of schooling: Shifting genetic influences across educational milestones in two Norwegian cohorts"

### Extended data figures for "Beyond years of schooling: Shifting genetic influences across educational milestones in two Norwegian cohorts"

**Extended Data Figure 1 | Educational attainment distribution in the MoBa cohort.**

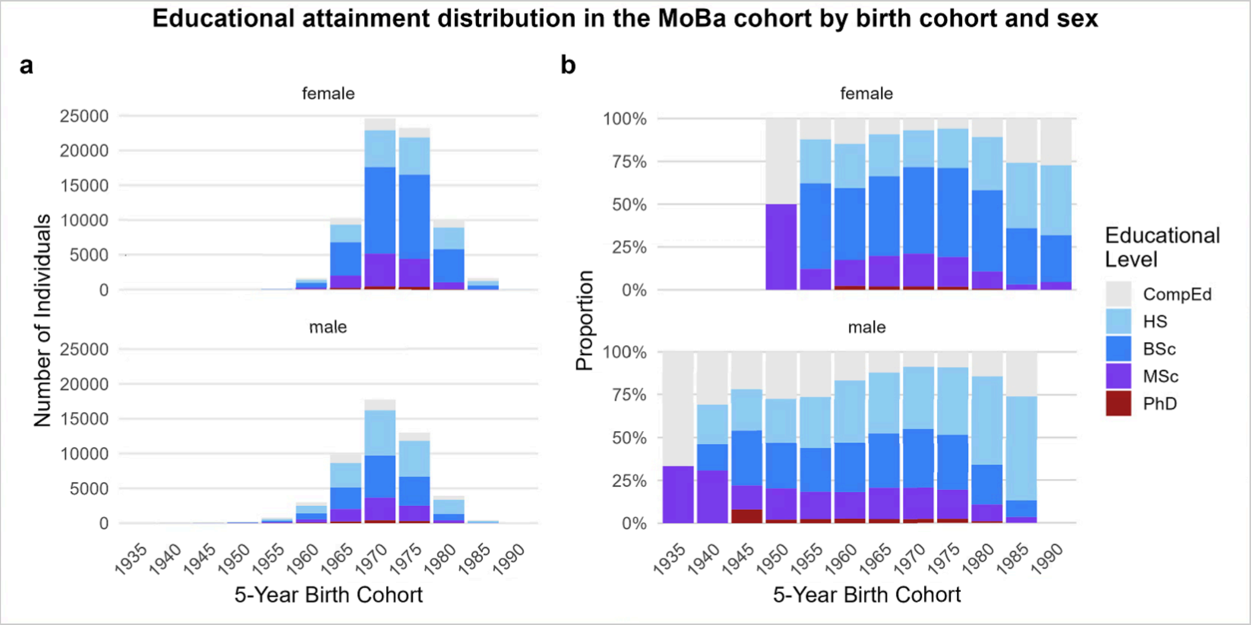


**Extended Data Figure 1 | EA across birth cohorts in the MoBa sample.** Distribution of EA in the Norwegian Mother, Father and Child Cohort Study (MoBa) sample (N = 120,527) stratified by sex and 5-year birth cohorts.

**a**, Absolute number of individuals achieving each educational level.

**b**, Proportion of individuals within each cohort achieving each educational level.

**Extended Data Figure 2 | Educational attainment distribution in the NTR cohort.**

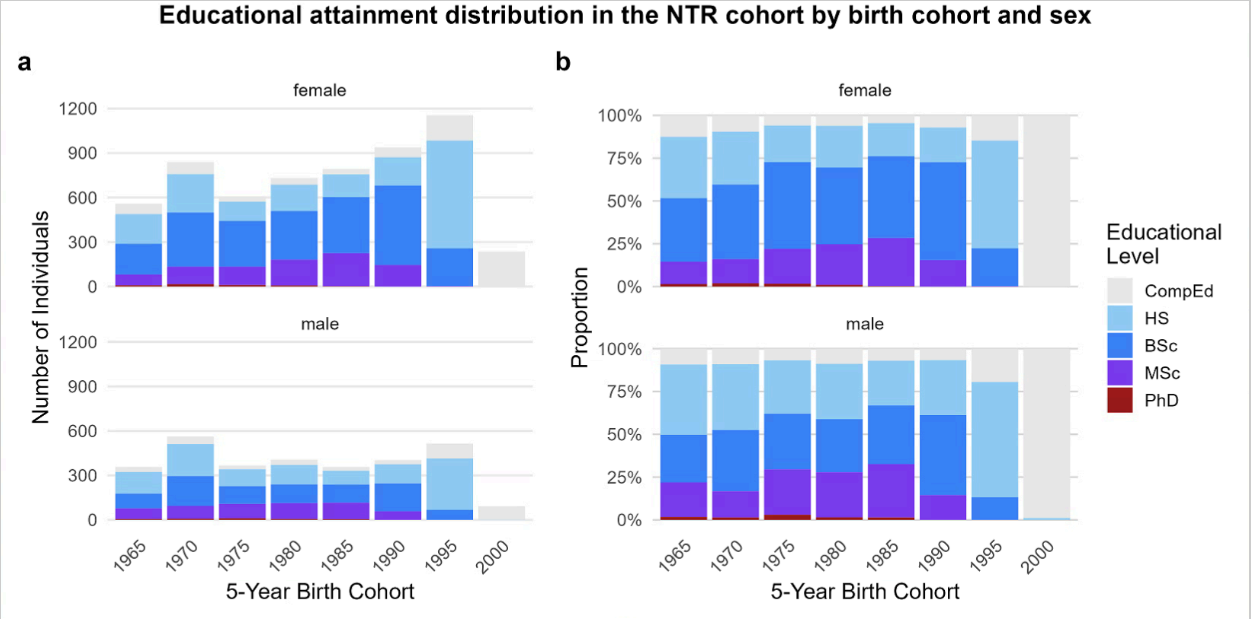


**Extended Data Figure 2 | EA across birth cohorts in the NTR sample.** Distribution of educational attainment in the Norwegian Twin Registry (NTR) sample (N = 8,910) stratified by sex and 5-year birth cohorts.

**a,** Absolute number of individuals achieving each educational level.

**b,** Proportion of individuals within each cohort achieving each educational level.

**Extended Data Table 1 | Mapping the Norwegian Standard Classification of Education (NUS2000) to analytical categories.**

| Tripartition of levels | Level | Level name | Class level | Included in Analyses | EA category | EduYears |
|---|---|---|---|---|---|---|
| | 0 | No education and pre-school education | Under school age | Excluded | | |
| Compulsory Education | 1 | Primary education | 1st - 7th class level | Included | CompEd | 7 |
| | 2 | Lower secondary education | 8th - 10th class level | Included | CompEd | 10 |
| Intermediate education | 3 | Upper secondary, basic | 11th - 12th class level | Included | CompEd | 12 |
| | 4 | Upper secondary, final year | 13th class level + | Included | HS | 13 |
| | 5 | Post-secondary not higher education | 14th class level + | Included | HS | 14 |
| Higher education | 6 | First stage of higher education, undergraduate level | 14th - 17th class level | Included | BSc | 16 |
| | 7 | First stage of higher education, graduate level | 18th - 19th class level | Included | MSc | 18 |
| | 8 | Second stage of higher education (postgraduate education) | 20th class level | Included | PhD | 21 |
| | 9 | Unspecified | | Excluded | | |

**Extended Data Table 1 | Mapping the Norwegian Standard Classification of Education (NUS2000) to analytical categories.**
The table details the official Norwegian Standard Classification of Education (NUS2000) levels. We mapped these levels to the five primary analytical categories used in this study: Compulsory Education (CompEd), High School (HS), Bachelor's (BSc), Master's (MSc), and Doctorate (PhD). The final column lists the conventional EduYears value assigned to each category. Educational levels 0 (pre-school) and 9 (unspecified) were excluded from all analyses. Adapted from Barrabés & Østli, 2016.
